# Resolving *Phytophthora* α2 hormone biosynthesis reveals species-specific mechanisms for mating type regulation

**DOI:** 10.64898/2026.04.23.720443

**Authors:** Milan Milenovic, Adam Jozwiak, Howard S. Judelson

## Abstract

Sexual reproduction by heterothallic species of *Phytophthora* involves communication between its two mating types by diterpenoid hormones derived from plant phytol. The genetics of mating type determination have remained unknown. We identified a conserved gene cluster that links mating identity to α2 hormone biosynthesis. Functional perturbation in *P. infestans* and pathway reconstruction in yeast showed that a cytochrome P450 in the cluster performs a mating type-specific step in hormone production. Comparative analyses revealed that the hormonal system is implemented through distinct genetic mechanisms in different species, including mating type-specific transcription in *P. infestans* and hemizygous inheritance in *P. capsici*. In self-fertile species the cluster is often eroded. These findings reveal a genetic basis for mating type and how conserved signaling systems can be evolutionarily remodeled.

## Introduction

Sexual reproduction in microbial eukaryotes is often gated by genetically defined mating types that control sexual development through diffusible signals (*1*). Although such systems are well characterized in filamentous fungi and yeasts, the basis of mating type communication remains poorly understood in non-model eukaryotes (*2*). Oomycetes, members of the stramenopile group alongside diatoms and brown algae, provide a useful system for understanding the diversity and evolution of mating systems as their sexual cycle differs markedly from those of fungi and even their close stramenopile relatives (*3*). Oomycetes are predominantly diploid and lack a free-living haploid phase; meiosis occurs in gametangia (oogonia and antheridia), where haploid nuclei fuse to form the zygote. Sexual reproduction begins with coordinated differentiation of gametangia in response to mating hormones.

Mating systems are particularly consequential in the oomycete genus *Phytophthora*, a group of diploid destructive plant pathogens in which sexual reproduction generates genetic diversity and durable survival spores (oospores) that complicate disease control worldwide (*4*). *Phytophthora* differs even from other oomycetes such as *Achlya*, where steroid hormones determine both sexual identity and mating compatibility. In *Phytophthora,* however, mating hormones are diterpenoids (Fig. 1A) rather than steroids, and mating type is not equivalent to sex. Each of the mating strains produces both oogonia and antheridia, making *Phytophthora* hermaphroditic.

**Fig. 1.**
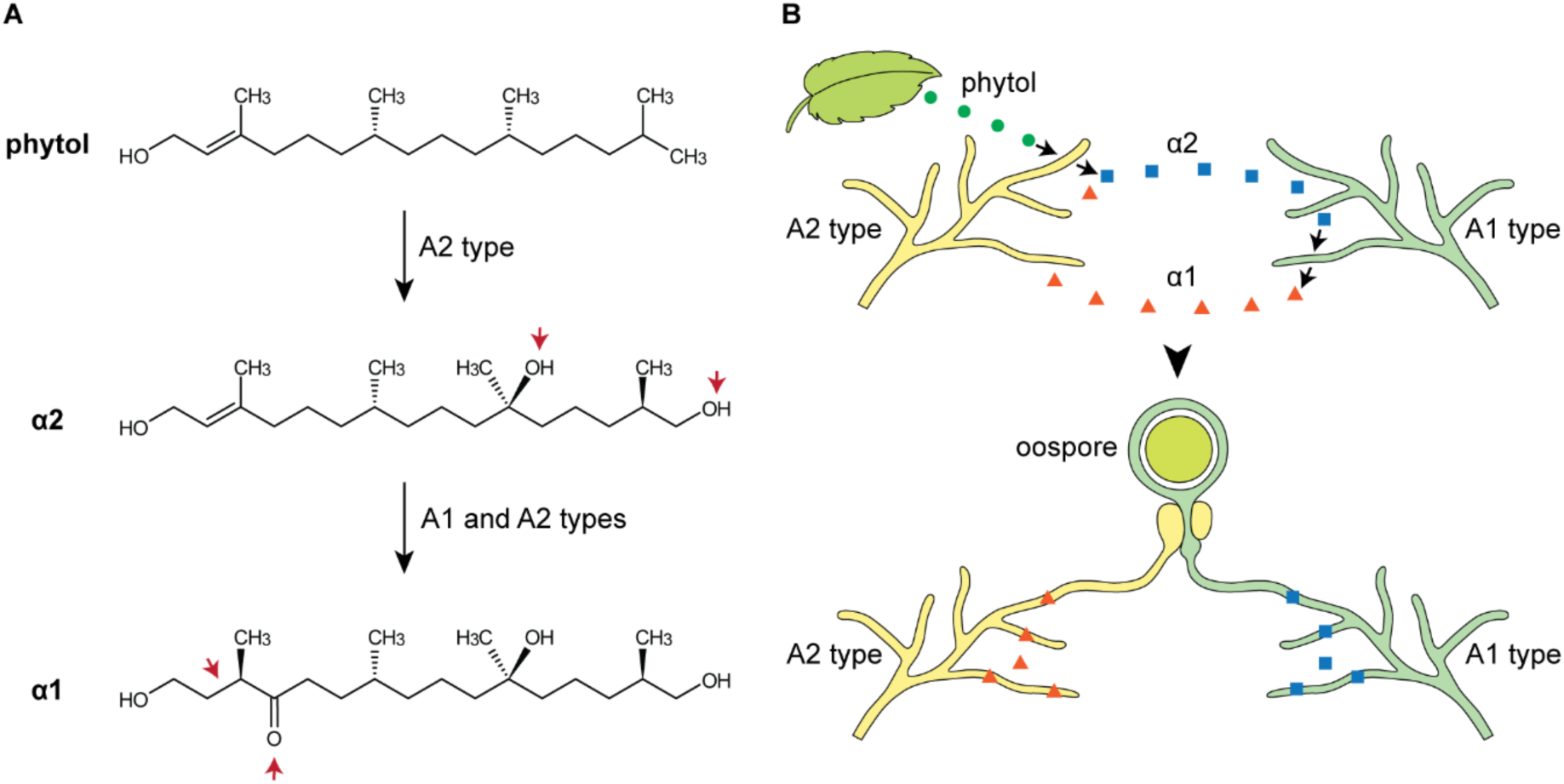
*Phytophthora* mating hormone biology. (A) Chemical structures of α1 and α2 mating hormones and their precursor phytol, a plant metabolite. Strains of A2 mating type convert phytol into α2. Both A1 and A2 mating types can further convert α2 into α1. (B) Illustration of hormone production and chemical communication between strains of opposing mating type. Hormone perception triggers sexual reproduction, resulting in durable, double-walled oospore. A1 strains are stimulated to form gametangia by α2, while A2 strains are stimulated by α1.

Some *Phytophthora* species lack mating types and are instead homothallic (self-fertile). Others are heterothallic, meaning that sexual reproduction can normally only occur between opposing mating types which are named A1 and A2 (*5*). In heterothallics, sexual development is triggered by two small molecules, known as the α1 and α2 hormones (Fig. 1) (*6*). Perception of these triggers the sexual cycle including gametangia development and the production of oospores. A1 strains are induced to undergo sexual development by α2 hormone, while A2 strains are induced by α1 (Fig. 1B) (*7*). The α2 hormone is produced by A2 strains from a plant-derived precursor, phytol (*6*, *8*). A1 strains, in turn, produce α1 by converting α2 which needs to be supplied by the A2. Interestingly, A2 strains are also capable of producing α1, without self-inducing sexual cycle (*6*, *8*, *9*). These hormones appear to be universal across heterothallic *Phytophthora* and detected even in some homothallic species, suggesting that the hormones are part of a mating system conserved within the genus (*6*). Understanding the asymmetry in hormone biosynthesis might point towards a genetic basis of mating type.

Biochemical studies have identified plausible intermediates in α2 production from phytol, consistent with involvement of oxidative enzymes, but the enzymes involved have remained unknown even two decades after α1 was characterized (*10*). Genetic studies in *Phytophthora* and relatives have not resolved the genetic basis of mating type, with studies in *P. infestans*, *Phytophthora capsici*, and their relative *Plasmopara viticola* (all members of the order Peronosporales) implicating different genomic regions as establishing mating type (*11–13*). Taken together, these results leave a possibility that mating type might be encoded in different ways in different species. Connecting the mating hormones to their biosynthetic pathway should help reveal how mating type is determined.

Here we make that connection. By combining comparative genomics across homo- and heterothallic Peronosporales with mating type-stratified expression data and functional tests, we identify a three-gene cluster associated with mating type that includes a cytochrome P450 monooxygenase (CYP) and two proteins that contain both START and FYVE domains, a combination not reported previously. We find that perturbing the P450 alters mating behavior and disrupts the production of phytol-derived metabolites including α2. We reconstruct α2 biosynthesis in yeast, demonstrating that two CYPs are responsible for production of the hormone. We find that this cluster and its association with mating-type is conserved across the Peronosporales, but the underlying genetics is not: in *P. infestans*, both mating types contain the cluster but an unlinked locus results in its A2-specific transcription. In contrast, in *P. capsici* the cluster is hemizygous in A2s and null in A1s, such that the chromosome transmitted from an A2 parent to A1 progeny lacks the cluster. Moreover, over 50% of homothallic species exhibit various forms of damage to the cluster, indicating that homothallism evolved from heterothallism. Together, these findings reveal how a conserved heterothallic signaling system can be built on shared pathway genes while evolving distinct mating type determinants.

## Results

### CYP repertoire and integrity in P. infestans

In *Phytophthora*, A2 strains synthesize α2 mating hormone via hydroxylation of two carbon atoms (*8*). Given that CYP enzymes commonly catalyze such hydroxylations, candidates for those enzymes were identified by mining a telomere-to-telomere *P. infestans* assembly using tBlastn against previously identified oomycete CYP proteins (*14*). This identified 27 loci for which membrane-anchor prediction, CYP-domain annotation, and structure modeling identified 17 encoding full-length CYP proteins with a putative N-terminal ER-anchor. The remaining 10 loci showed frameshifts or premature stop codons relative to orthologs in other species. To allow interspecific comparisons of the functional CYPs, we also mined 114 Peronosporales genomes for putative CYPs which OrthoFinder clustered into 30 orthogroups (Fig. S1 and Table S1). Phylogenies within each orthogroup were largely congruent with the species tree (Fig. S2). This approach ensured a comprehensive CYP repertoire across species, allowing inclusion of unannotated genomes in our subsequent comparative analyses.

### One CYP shows mating type-associated expression

We hypothesized that a CYP involved in α2 production would show mating type-specific expression. To test this, we performed differential gene expression analysis on eleven A1 and nine A2 *P. infestans* strains consisting of 13 geographically diverse field isolates and 7 F1 progeny of a cross (1306 × 618, strains beginning with a letter “M” in Fig. 2). Out of 17 CYP genes, 14 were expressed at similar levels in both mating types, while two were not detectably expressed in either. One (PITG_07424) was expressed at significantly higher levels (≥90-fold) in A2 strains compared to A1s (FDR=8.94 × 10^-5^), with normalized CPMs ranging from 155 to 2884. In contrast, PITG_07424 showed low to no expression in all A1 strains, with CPM values ranging from 0 to 14 (Fig. 2). In all A1 strains with non-zero expression, reads consistently mapped only to the 3’ end of PITG_07424 (Fig. S3), whereas coverage across other CYP genes was uniform. A2-specific expression was confirmed by RT-qPCR of A2 strain 618 and A1 strain 1306. Analysis of RNA-seq data from 32 additional F1 progeny of the A1 mating type using JBrowse2 also found that PITG_07424 is not expressed at significant levels in that mating type.

**Fig. 2.**
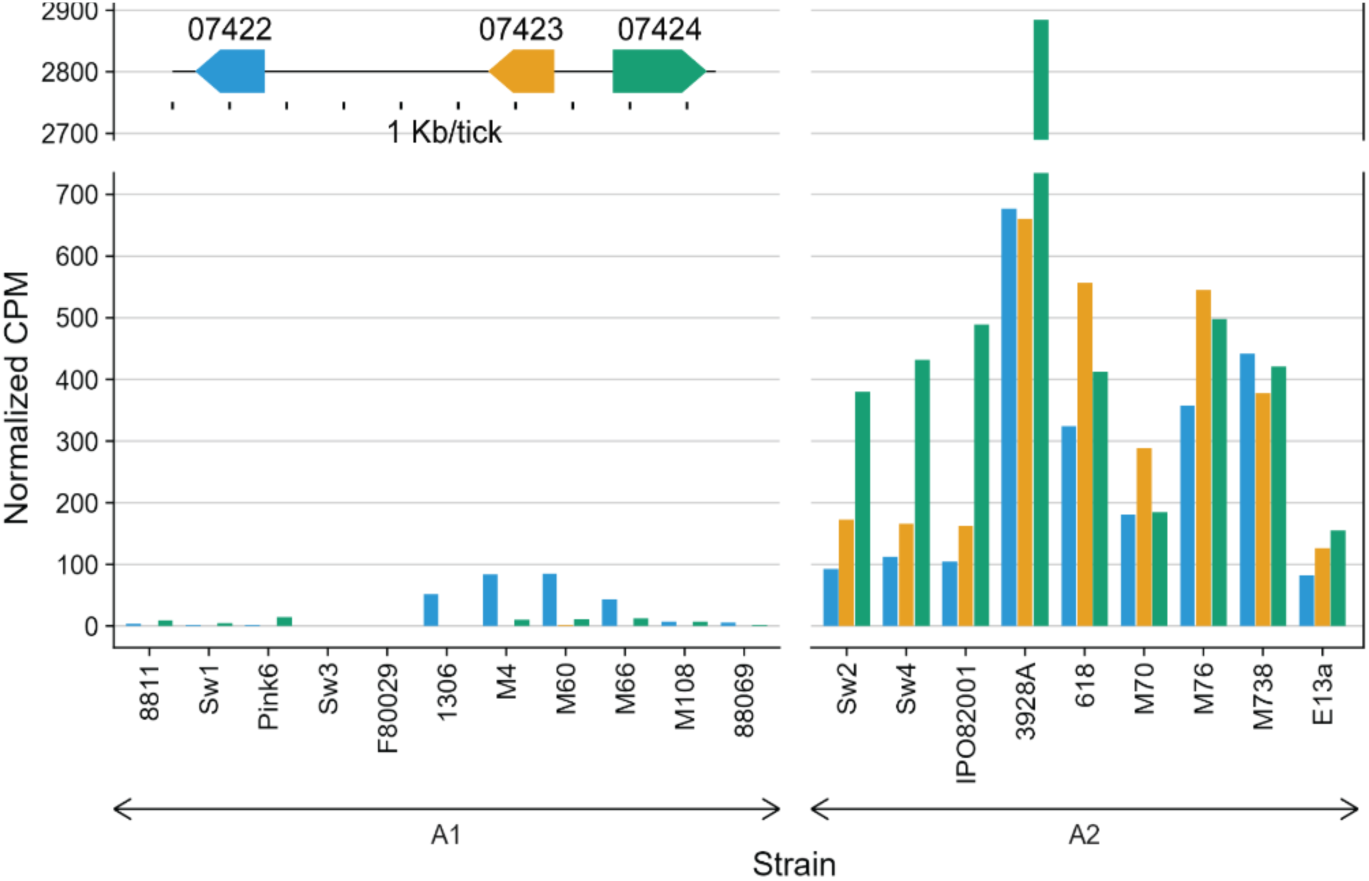
Differential expression of a three-gene locus between A1 and A2 mating type. Normalized expression levels of genes PITG_07422 (blue), PITG_07423 (orange), and PITG_07424 (green) are shown for each A1 and A2 strain. The inset schematic (top left) shows gene orientation, lengths, and relative genomic positions within the cluster; the same gene order was observed in each strain.

Differential expression analysis also revealed two additional A2-specific genes, which form a cluster with PITG_07424 (Fig. 2). These were PITG_07422 and PITG_07423 which exhibited 28-fold and 1900-fold higher levels of expression in the A2, respectively. Much like the CYP (PITG_07424), the expression of PITG_07423 was perfectly correlated with A2 mating type. PITG_07422, however, showed minor levels of expression in some A1 strains. PITG_07422 and PITG_07423 encode similar proteins, both containing putative START and FYVE domains which will be addressed in more detail below. No other unannotated or annotated regions of the *P. infestans* genome exhibited mating type-biased expression. Thus, the analysis identified a single candidate mating type locus to be functionally tested.

### Altering expression of CYP PITG_07424 switches mating phenotype

When an A2 strain encounters an A1, normally oospores are generated while asexual sporulation is suppressed (Fig. 3Ai). This changed when both alleles of PITG_07424 were knocked out in A2 strain 618 using the CRISPR/Cas12a system. The resulting mutants (ι107424) failed to produce oospores when paired with A1 strains but continued to produce asexual sporangia (Fig. 3Aii). In addition, the mutants now produced oospores when paired with either of several A2 strains (Fig. 3Aiii). The resulting interaction zones also lacked aerial hyphae and asexual spores, as is the case in normal matings.

**Fig. 3.**
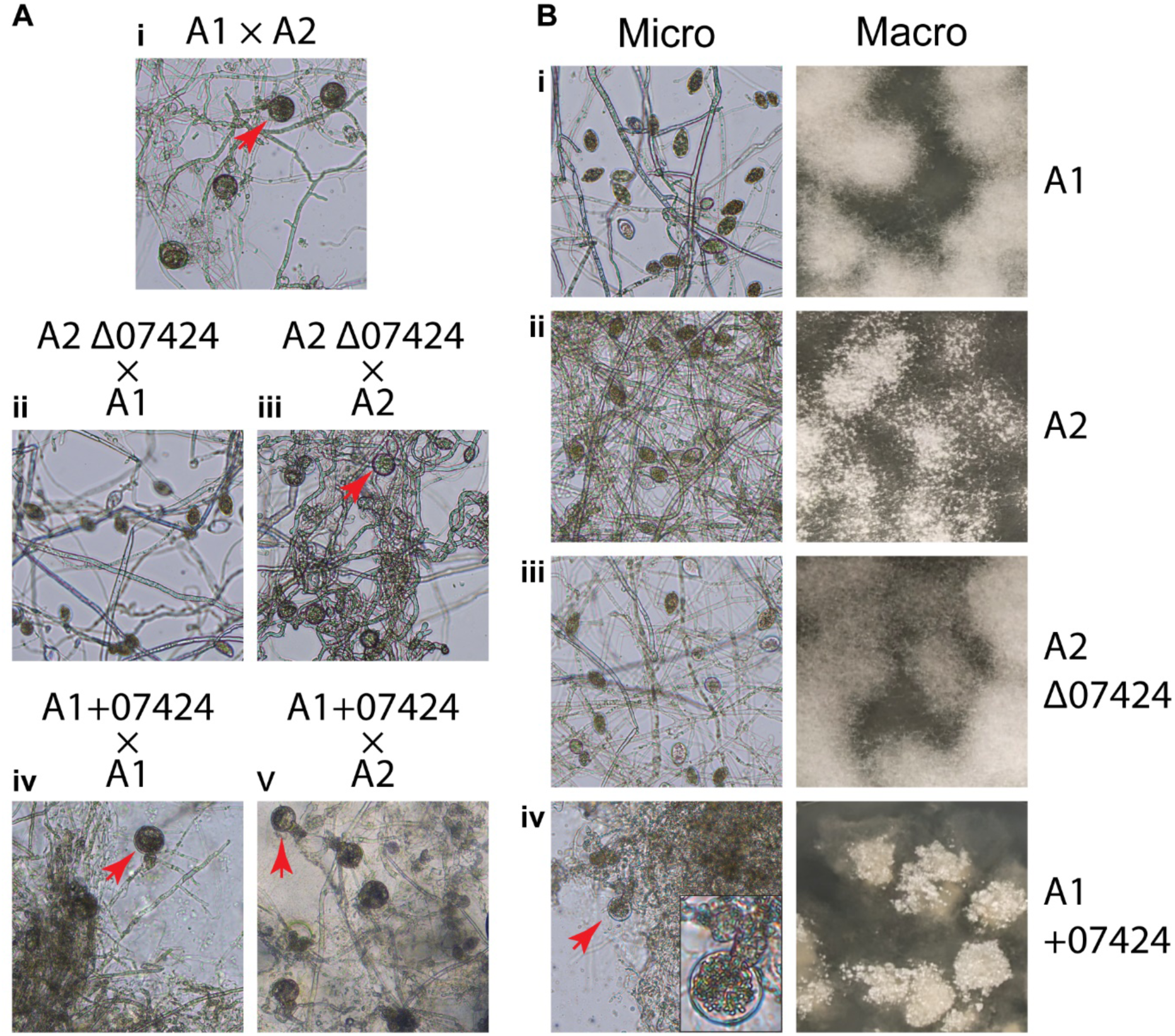
Phenotypes of *P. infestans* wild-type and engineered strains. (A) Light microscopy of co-cultures with the indicated strain combinations. Asexual spores (sporangia) are smaller, ellipsoidal structures while the sexual spores (oospores) are larger spherical structures. One oospore is marked by a red arrow in each image with oospores. (B) Light microscopy (left), and macroscopic image (right) of the same strains grown in single culture. Microscopy cultures were grown on rye agar, while macroscopic images are of mycelia on the surface of rye broth, both supplemented with phytol.

The disruption of PITG_07424 in the A2 strain also resulted in the loss of the “lumpy” colony morphology typical of A2 strains of *P. infestans.* Although its biological significance is unknown, this is due to rapid hyphal branching which results in lumps on the top of the hyphal mat (Fig. 3Bii). This is not seen in A1 strains (Fig. 3Bi). However, deletion of PITG_07424 in the A2 resulted in a fluffy A1-like morphology (Fig. 3Biii).

Phenotype changes were also observed when PITG_07424 was expressed ectopically in an A1 isolate using a strong constitutive promoter. Cultures of the resulting A1+07424 strain alone developed oospores (Fig. 3Biv). This can be compared to single cultures of an A1 (Fig. 3Bi) or A2 (Fig. 3Bii) which only produce the lemon-shaped asexual sporangia. Moreover, the A1 strain expressing PITG_07424 acquired the lumpy phenotype (Fig. 3Biv). Oospores were also seen when the A1+07424 strain was paired with either wild-type A1 or A2 strains (Fig. 3Aiv and v). Together, the overexpression and knockout experiments confirmed the involvement of CYP PITG_07424 in mating type determination.

### Altering expression of CYP PITG_07424 also affects hormone production

To confirm our hypothesis that expression of the CYP determines α2 production, we performed untargeted and targeted LC-MS analyses on wild-type *P. infestans* and the transformants. A1 and A2 strains displayed distinct phytol-derived metabolite profiles (Fig. 4). In wild-type A2 strains, peaks matching the masses of the previously reported mating hormones α1 and α2 were detected. These were not detected in cultures grown without phytol. As expected, the hormones were not detected in wild-type A1 as α2, the precursor to α1, was not provided (*8*). Next, we looked at the modified strains. The Δ07424 A2 strain lacked detectable A2-specific metabolites (including α1 and α2, indicated by arrows in Fig. 4) and instead its profile resembled that of the A1. By contrast, the predominant peak in the A1 strain expressing PITG_07424 (A1+07424) was α1. A very small peak matching α2 was present. Our interpretation of this is that the transformant was able to produce α2, which was then mostly converted to α1.

**Fig. 4.**
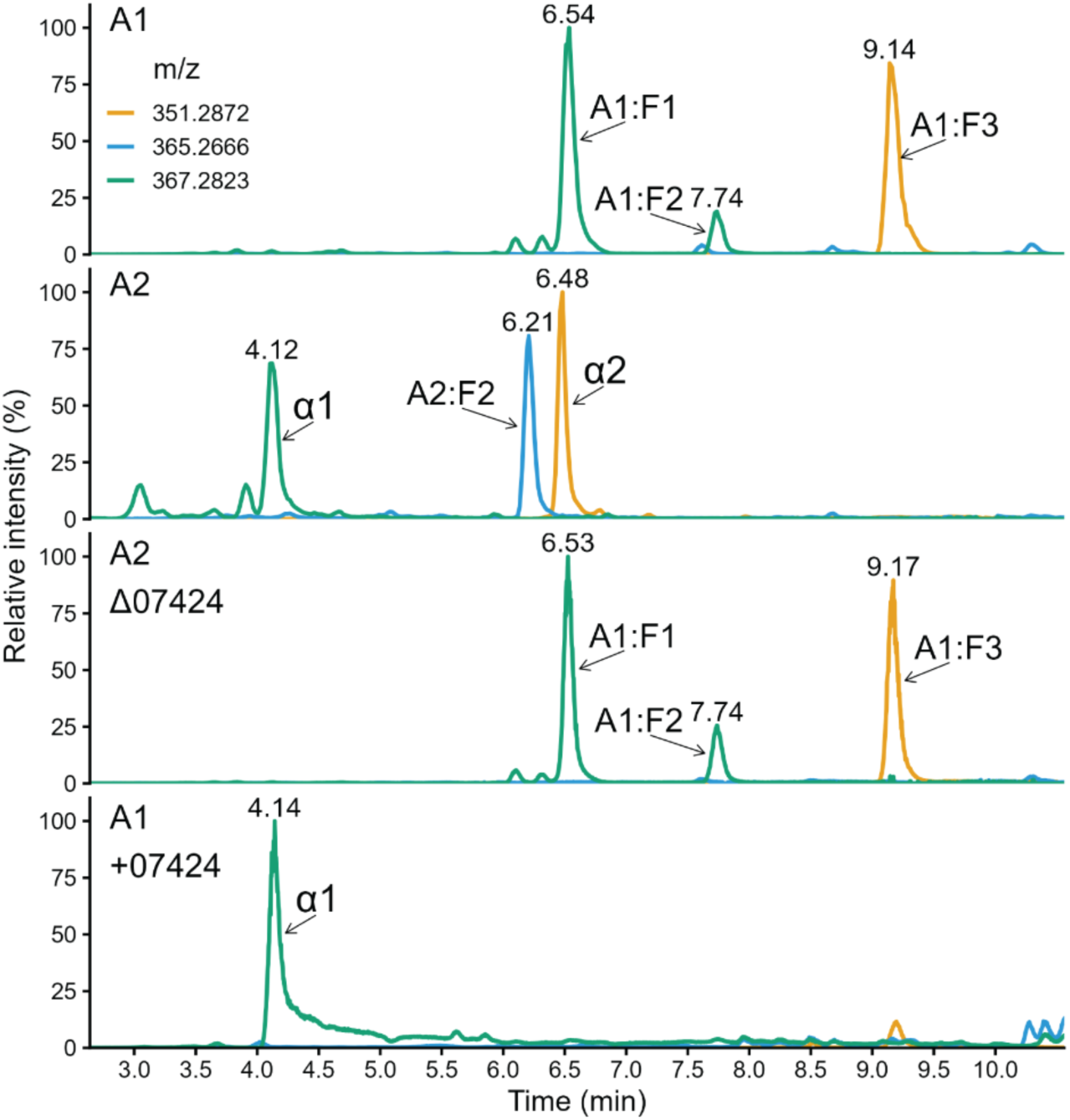
Extracted ion chromatograms (EICs) showing major phytol-derived metabolites in A1 and A2 wild-type strains, an A2 Δ07424 biallelic knockout, and an A1 strain overexpressing 07424 (A1+07424). Retention times in min are indicated above peaks; arrows denote peaks with annotated compounds including the mating hormones, α1 and α2. Traces correspond to the extracted masses: m/z 351.2871 (orange), 365.2666 (blue), and 357.2823 (green). Similar results were obtained from mycelia extracts.

Although prior studies of the *Phytophthora* mating hormones have focused on α1 and α2 alone, our untargeted analysis of wild-type strains also discovered additional phytol metabolites distinct from the known mating hormones. Several of these are labelled in Fig. 4 (A2:F2, A1:F1, A1:F2, and A1:F3). One novel A1 metabolite had biological activity and is discussed in later sections.

We confirmed the identity of α1 and α2 in our analyses, and partially characterized the novel metabolites, by derivatization of A1 and A2 extracts using Girard’s T reagent. In A2 extracts, the α1 peak was diminished by the reaction, consistent with the presence of a carbonyl (keto) functional group in this metabolite, whereas α2 and A2:F2 remained unchanged (Fig. S4). In A1, all phytol metabolites showed a reduction, also consistent with the presence of a keto group.

Experiments using deuterium-labelled phytol (phytol-*d*6) also showed that α1 and all three A1 phytol metabolites lost two deuterium atoms, while α2 and A2:F2 were still heavier by six deuterium atoms (Table S2). Introduction of a keto group at C4 (ω-end), as observed for the α1 hormone and A1 metabolites, resulted in the loss of two deuterium atoms, yielding a net mass increase of +4 Da relative to unlabeled phytol-derived metabolites. A mass shift of +6 relative to unlabeled counterparts corresponds to oxidation at α-end of the molecule, as all deuterium atoms remained. This metabolomics analysis identified PITG_07424 as a key determinant of the A2 metabolome, including α2, paving the way for pathway reconstruction.

### Reconstruction of α2 biosynthetic pathway in yeast

To test whether PITG_07424 catalyzes the initial step in converting phytol to α2, or a later step, we expressed that protein in *S. cerevisiae* WAT11 grown in media supplemented with phytol. WAT11 expresses a plant NADPH-P450 reductase (CPR) which can enhance the catalytic efficiency of heterologous CYPs. LC-MS analysis detected no phytol-derived metabolites in PITG_07424-expressing *S. cerevisiae* WAT11 or an empty-vector control. Similar results were obtained with a FLAG-tagged version of PITG_07424, in which its expression was confirmed by western blotting. Co-expressing PITG_07424 with the *P. infestans* CPR also yielded no detectable phytol-derived products in either cells or medium. These results suggest that PITG_07424 does not catalyze the initial hydroxylation of phytol but instead acts on a hydroxylated intermediate produced by another CYP.

To test this, we screened the other 16 CYP genes from *P. infestans* using the same yeast system and found that only PITG_21524 and PITG_21759 could metabolize phytol. PITG_21524 produced a hydroxyphytol that matched a metabolite in *Phytophthora* extracts (peak 2 in Fig. 5A; RT=11.7 min, m/z=335.2920, [M+Na]^+^, identical MS^2^). An additional peak corresponding to dihydroxyphytol was detected (RT=8.51 min, m/z=351.2870, [M+Na]^+^). This peak was not detected in *Phytophthora* extracts and likely reflects further metabolism of hydroxyphytol by endogenous yeast enzymes. The other gene, PITG_21759, metabolized phytol into a different hydroxyphytol (RT=12.16 min, m/z=295.2995, [M-H_2_O+H]^+^), which was also detected in *P. infestans* cultures (Fig. S5).

**Fig. 5.**
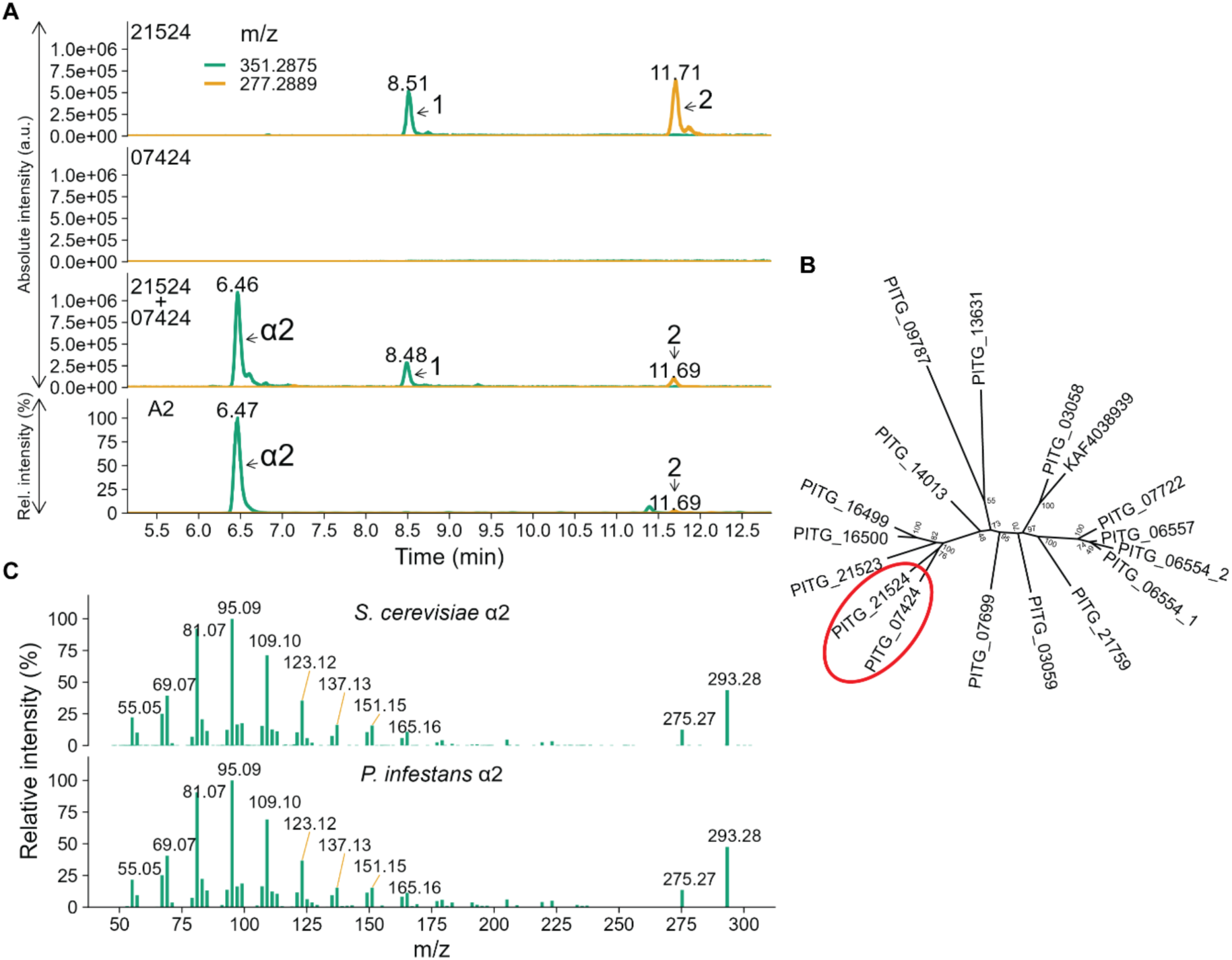
Reconstruction of the α2 biosynthetic activity in yeast. (A) EICs from *Saccharomyces cerevisiae* expressing *Phytophthora infestans* cytochrome P450 (CYP) genes 21524, 07424, or both (21524+07424), showing production of phytol-derived metabolites. For comparison, the *P. infestans* A2 culture EIC is shown below. Phytol was added to all cultures. Traces are shown for m/z 351.2875 (green; putative dihydroxyphytol observed as [M+Na]^+^) and 277.2889 (orange; putative hydroxyphytol observed as [M-2H_2_O+H]^+^). Retention times (min) are indicated above peaks and arrows label peak identities. Yeast EIC panels use identical absolute intensity scaling; the *P. infestans* A2 EIC is displayed on a relative intensity scale. (B) Phylogenetic relationship of *P. infestans* CYP genes; 07424 and 21524 are highlighted (red). Numbers at each node show bootstrap value (%). (C) MS^2^ spectra of α2 produced in yeast (top) and in *P. infestans* A2 cultures (bottom).

To test if any of these hydroxyphytols are α2 precursors, we co-expressed PITG_07424 with PITG_21524 or PITG_21759. Only PITG_07424 acted on the PITG_21524-derived hydroxyphytol, producing a metabolite with the same accurate mass, retention time, and MS^2^ spectrum as α2 in *P. infestans* extracts (Fig. 5A, C). The amounts of the original hydroxyphytol intermediate (peak 2) and the yeast-specific dihydroxyphytol (peak 1) were markedly reduced. Injection of yeast-produced metabolite, spiked with purified *P. infestans* α2, yielded a single chromatographic peak with a matching, single MS^2^ spectrum (Fig. S6). Consistent with their partnership, phylogenetic analysis revealed that PITG_07424 and PITG_21524 form a well-supported clade distinct from other CYPs in the genome (Fig. 5B). However, PITG_21524 is expressed in both mating types and is located on a different chromosome (Chr 3) than PITG_07424 (Chr 13). Thus, reconstructing the α2 pathway in yeast revealed that it occurs in two steps involving two related CYPs: PITG_21524 as the first required phytol monooxygenase and PITG_07424 as the hydroxyphytol monooxygenase.

### Analysis of novel phytol metabolites reveals a new bioactivity

Untargeted statistical metabolomics of media extracts from A1 and A2 strains, with and without added phytol, revealed additional phytol-derived metabolites beyond the known mating hormones (Fig. 4). The major A1-derived compounds (A1:F1 and A1:F3) had comparable peak areas to α1 and α2 from A2 strains and the same nominal mass as α1 or α2 but longer retention times (6.54 versus 4.12, and 9.14 versus 6.48) and distinct MS^2^ spectra (Fig. S7). Both mating types produced multiple additional phytol-derived metabolites in lower but easily detectable amounts (>20 in total). Incorporation of deuterium-labeled phytol caused the expected mass shifts in these metabolites, confirming that they are derived from phytol.

Intriguingly, while A1 cultures had no response to A1:F3, A2 hyphae displayed accelerated growth towards it which was observable in less than 24 h (Fig. 6B). The zone growing towards A1:F3 was less dense, had reduced aerial hyphae, and fewer asexual sporangia compared to normal cultures. No oospores were observed. The suppression of asexual sporulation persisted for at least 10 days; normally asexual spores would appear by three days. Thus, our investigation of other phytol metabolites revealed that mating signaling extends beyond canonical hormones, indicating a more complex chemical communication system.

**Fig. 6.**
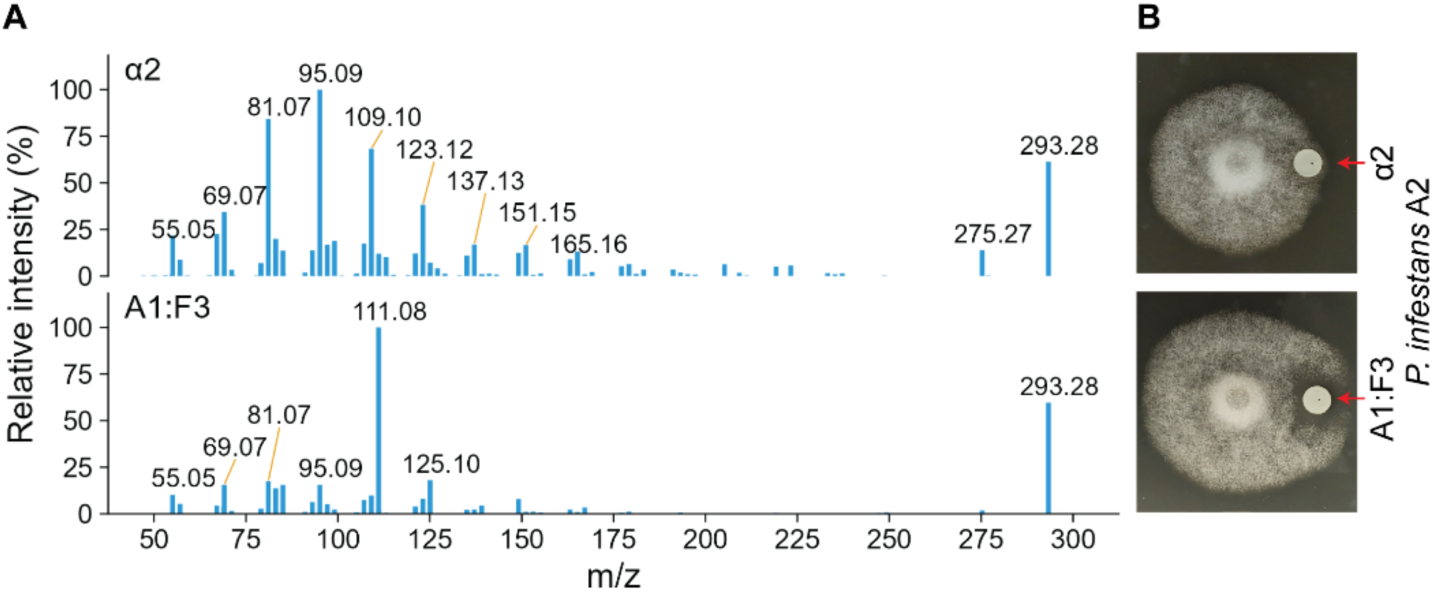
A novel bioactive phytol-derived metabolite produced by A1. (A) MS^2^ spectra of the *P. infestans* α2 hormone (top) and a distinct phytol-derived metabolite produced by *P. infestans* A1 (A1:F3; bottom). (B) Images show paper-disc diffusion bioassay showing the growth response of *P. infestans* A2 to purified α2 (top) and A1:F3 fractions (bottom).

Although A1:F3 has the same monoisotopic mass as α2, it has a distinct retention time (Fig. 4) and a clearly different MS2 spectrum (Fig. 6A). Notably, its fragmentation pattern is more like that of α1 than α2: metabolite A1:F3 and α1 share the same major product ions, consistent with fragmentation from the α-end of the molecule containing a keto group and lacking the double bond characteristic of α1 (Fig. S8). This pattern, together with the known biosynthetic logic of the pathway, suggests that A1:F3 is not an α2-like compound, but rather an α1-like derivative. Specifically, our data support a model in which A1 strains can oxidize phytol by introducing one hydroxyl group, removing the double bond, and forming the keto group, but cannot install the second hydroxyl group, a step attributed to the A2-specific CYP (PITG_07424) activity.

Consistent with this interpretation, feeding experiments with phytol-*d*6 showed that A1 produced a labeled A1:F3 species carrying only four deuterium atoms, indicating loss of two deuterium atoms at the position oxidized to the keto group (Fig. S9). MS^2^ spectrum of the labelled A1:F3 also agreed with this interpretation (Fig. S9). Taken together, these observations strongly support the conclusion that A1:F3 is best explained as an α1-like product that lacks the A2-dependent hydroxylation step.

We considered the possibility that some of the novel peaks evident in Fig. 4 might represent biosynthetic intermediates of the mating hormones rather than phytol-derived branch products *per se*. To test this, we collected and analyzed the metabolite-carrying paper disks at the end of the bioassay experiment in which the mating hormones and four novel compounds had been tested (Fig. 4). LC/MS showed that most applied metabolites were not further metabolized, with only trace amounts of more oxidized derivatives detected. The exception was the α2 mating hormone, which was almost completely converted to α1 by A1 strains, whereas A2 strains converted a smaller but detectable fraction of added α2 to α1 (Table S2). These observations are consistent with the previously published model in which A1 strains rely on exogenous α2 to produce α1 and prior evidence that A1 strains cannot synthesize α1 directly from phytol.

### Promoters within the gene cluster are functional but not active in A1 strains

Comparisons of A1 and A2 strains indicated that the A2-specific expression of the gene cluster is not determined by polymorphisms within their regulatory sequences. While examinations of the 973-nt promoter region shared by PITG_07424 and PITG_07423 in A1 strains 1306 and 6629 and A2 strains 550 and 618 identified one SNP and one small indel, these were not mating type-specific (Fig. 7A). Moreover, segregation of these eight haplotypes (identified from phased assemblies) in 108 F1 progeny from 1306×618 and 550×6629 crosses showed no association with mating type. While phylogenetic footprinting across the Peronosporales as well as motif discovery using MEME identified several conserved motifs in their promoter regions (including three upstream conserved elements at -243, -315, and -599 bp; Fig. S10), these did not include the SNPs detected between the strains. A lack of correlation with mating type was also observed for SNPs in the coding sequence and 3’ untranslated region of PITG_07424.

**Fig. 7.**
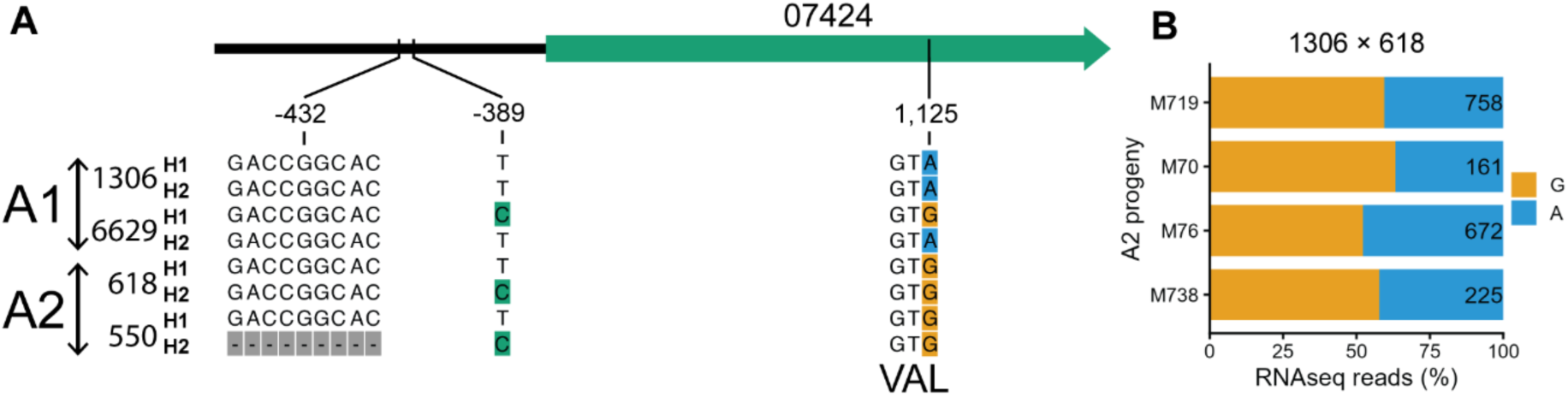
Promoter polymorphisms are not associated with mating type-specific expression of PITG_07424. (A) Schematic and sequence comparison of promoter and coding-sequence (CDS) variants of PITG_07424 across two A1 and two A2 strains. Coordinates are shown relative to the PITG_07424 start codon. The CDS polymorphism is shown in-frame; the substitution is synonymous (both codons encode valine (Val)). (B) Allele-specific read counts at SNP position 1,125 in RNA-seq data from 1306 × 618 F1 progeny (A2). Bars show the proportion of reads supporting the A versus G allele at the focal site; numbers within bars indicate total read coverage at that position (A+G reads).

Direct evidence for the functionality of the quiescent alleles in A1 strains was yielded by allele-specific RNA-seq analysis using a SNP within the coding sequence of PITG_07424, and SNPs flanking the gene to follow their segregation. Both silent alleles in the A1 strains could be transmitted to and expressed in A2 progeny (Fig. 7B). It follows that mating type is regulated by a locus unlinked to the gene cluster.

Also consistent with the involvement of an unlinked locus were the results of an attempted ectopic expression experiment using PITG_07424. When the gene plus native (896 nt) promoter from an A2 strain was transformed into an A1 strain, no expression was detected by RT-PCR among 10 transformants, in contrast to 8 out of 10 transformants expressing the gene when driven by constitutive promoter. No phenotypic or metabolic change was observed either. This, combined with differentially expressed region analysis, suggests that A1 strains may not have a required transcription factor or chromatin modifier that affects transcription of the cluster genes.

Due to the possible involvement of a chromatin modifier (*15*), we considered whether histone deacetylation might block expression in A1 strains. However, the histone deacetylase inhibitor trichostatin A did not restore PITG_07424 expression in the A1 based on RT-qPCR. Trichostatin also did not cause any mating hormone to be produced in the A1 based on LC-MS. Metabolomics showed that TSA broadly reduced metabolite abundances in both mating types which indicates that trichostatin was affecting *P. infestans* (Fig. S11 and Fig. S12). However, α2 and two unknown metabolites increased in abundance (Fig. S13).

### Cytochrome PITG_07424 is part of a cluster that also encodes START-FYVE proteins

As noted earlier, PITG_07424 is clustered with two other genes, PITG_07423 and PITG_07422, that also exhibit A2-specific expression (Fig. 2). The cluster is highly conserved in the Peronosporales based on the analysis of 114 genomes (Fig. S2 and Table S3). However, no orthologs of any of the three genes were identified outside the Peronosporales (genera *Halophytophthora*, *Phytopythium*, *Pythium,* and more distant order Saprolegniales) (Fig. S2).

Conservation of the cluster within six representative heterothallic species is illustrated in Fig. 8 with a complete dataset represented in Fig. S2 and Table S3. Synteny was incomplete outside of the three-gene cluster. All heterothallic species also contained an intact ortholog of the second CYP needed to make α2, PITG_21524, but elsewhere in the genome.

**Fig. 8.**
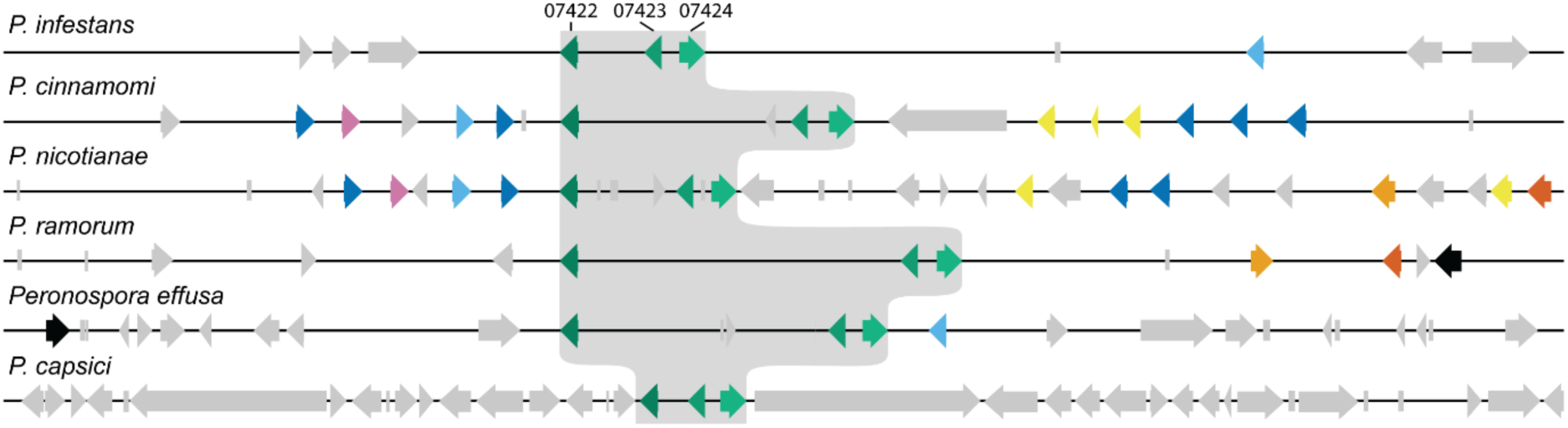
Synteny of genes flanking the mating gene cluster. Gene models are shown across a 93 Kb genomic interval adjacent to the cluster in six Peronosporales species having annotated genomes. Orthologous genes are shown in the same color. Within the mating gene cluster (gray shaded area), this includes PITG_07422, PITG_07423, and PITG_07424 which are represented by shades of green (from dark to light, respectively). Genes not conserved between the species (non-syntenic) are in light gray. Orthology was tested by BLASTP matches to the *Phytophthora infestans* proteome (E < 1×10^-3^; query coverage ≥85%; identity >55%) or by all to all blastp for genes without a hit in *P. infestans*.

In contrast with the heterothallic Peronosporales, over 50% (32 species) of homothallics exhibited disruption or loss of at least one of the three genes. This is illustrated in Fig. 9 for the homothallics *P. versiformis*, *P. boehmeriae*, and *P. kernoviae*. Six species of *Phytophthora* clade 2 lack orthologs or remnants of any of the three genes in the reference genomes analyzed.

**Fig. 9.**
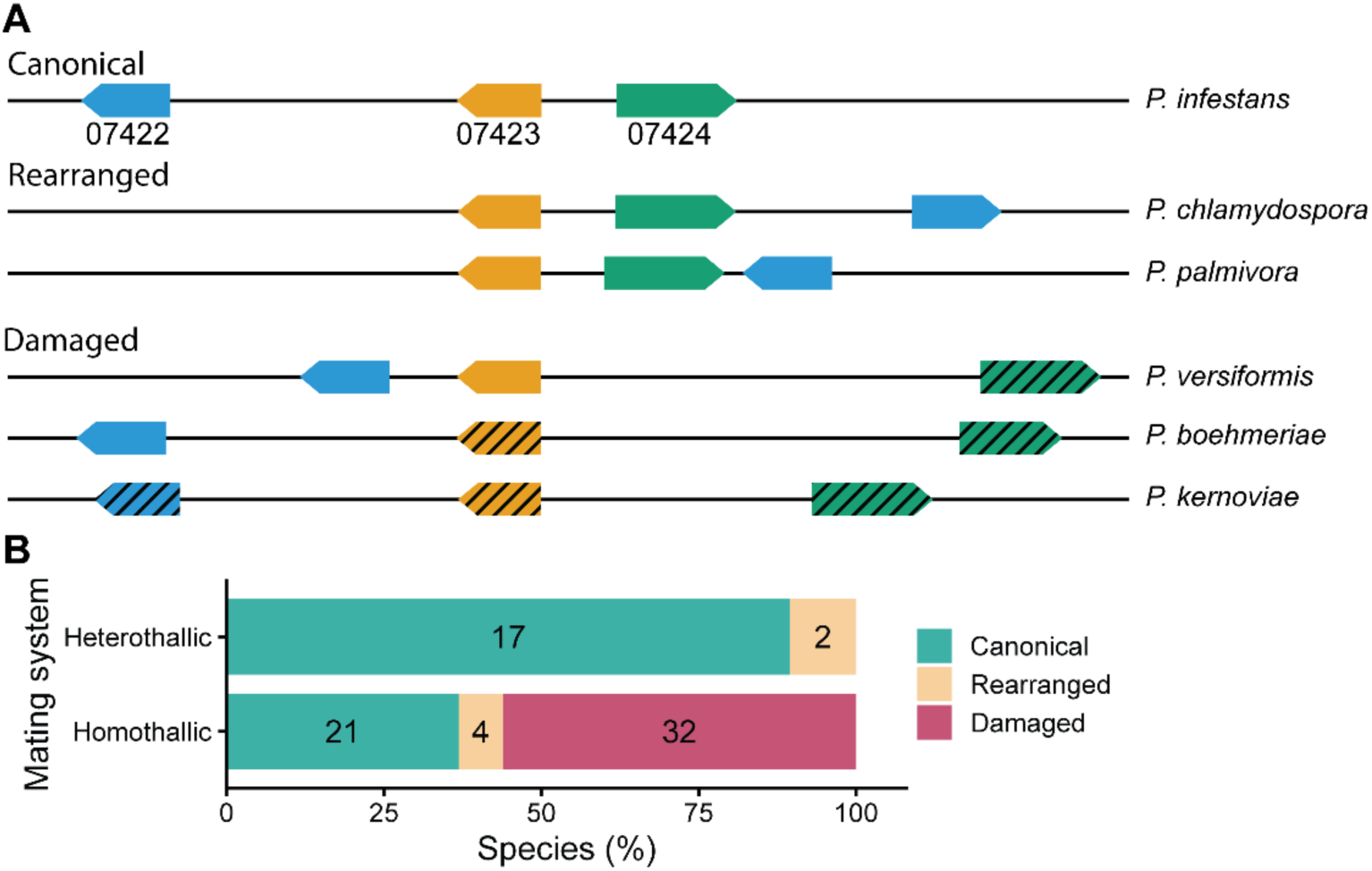
Integrity of the three-gene cluster across Peronosporales and association with reproductive mode. (A) Representative genomic configurations of 07422, 07423, and 07424 orthologs across Peronosporales, showing gene order, orientation, and intergenic distances. Damaged genes are indicated by hatched fill. Canonical and rearranged example species are heterothallic; example species for damaged genes are homothallic. (B) Proportion (and number) of species exhibiting an intact cluster, rearrangement, or damage, stratified by heterothallic versus homothallic reproductive mode. Species were classified as “damaged” if at least one of the three orthologs was disrupted.

Since genes PITG_07423 and PITG_07422 are tightly associated with mating type, their potential functions were studied in more detail. Both encode proteins with an N-terminal START domain followed by a C-terminal FYVE-like domain (Fig. S14 and Fig. S15). Matches to these domains in the NCBI Conserved Domain Database were weak but were detected for many of the *Phytophthora* orthologs.

### Deletion of the cluster from one haplotype determines mating type in P. capsici

As in *P. infestans,* RNA analysis indicated that the gene cluster was also expressed only in A2 isolates of *P. capsici.* However, instead of the case in *P. infestans* where the cluster was in both mating types but differentially regulated, a structural difference exists within *P. capsici*: the cluster is absent from A1s while hemizygous in A2s. This was initially detected by comparing long-read data from an A1 and an A2 strain (Fig. 10B) and later corroborated using Illumina reads from four strains of known mating type in NCBI SRA (A1: BYA5, 55330; A2: LT263, LT1534). As shown in the A2 track in Fig. 10B, only about half of the long reads included the gene cluster, while in the A1 track all reads included the deletion. Additional genome assemblies lacking corresponding read data further supported this observation: three A1 genomes (strains 05-06, KPC-7, and JHAI-7) lacked the cluster, whereas the A2 genome (MY-1) contained it. The deletion encompasses the CYP and two START–FYVE genes, leaving flanking genes intact (Fig. 10A,C). BLAST analysis failed to detect the three genes elsewhere in the genome, excluding the possibility of a translocation.

**Fig. 10.**
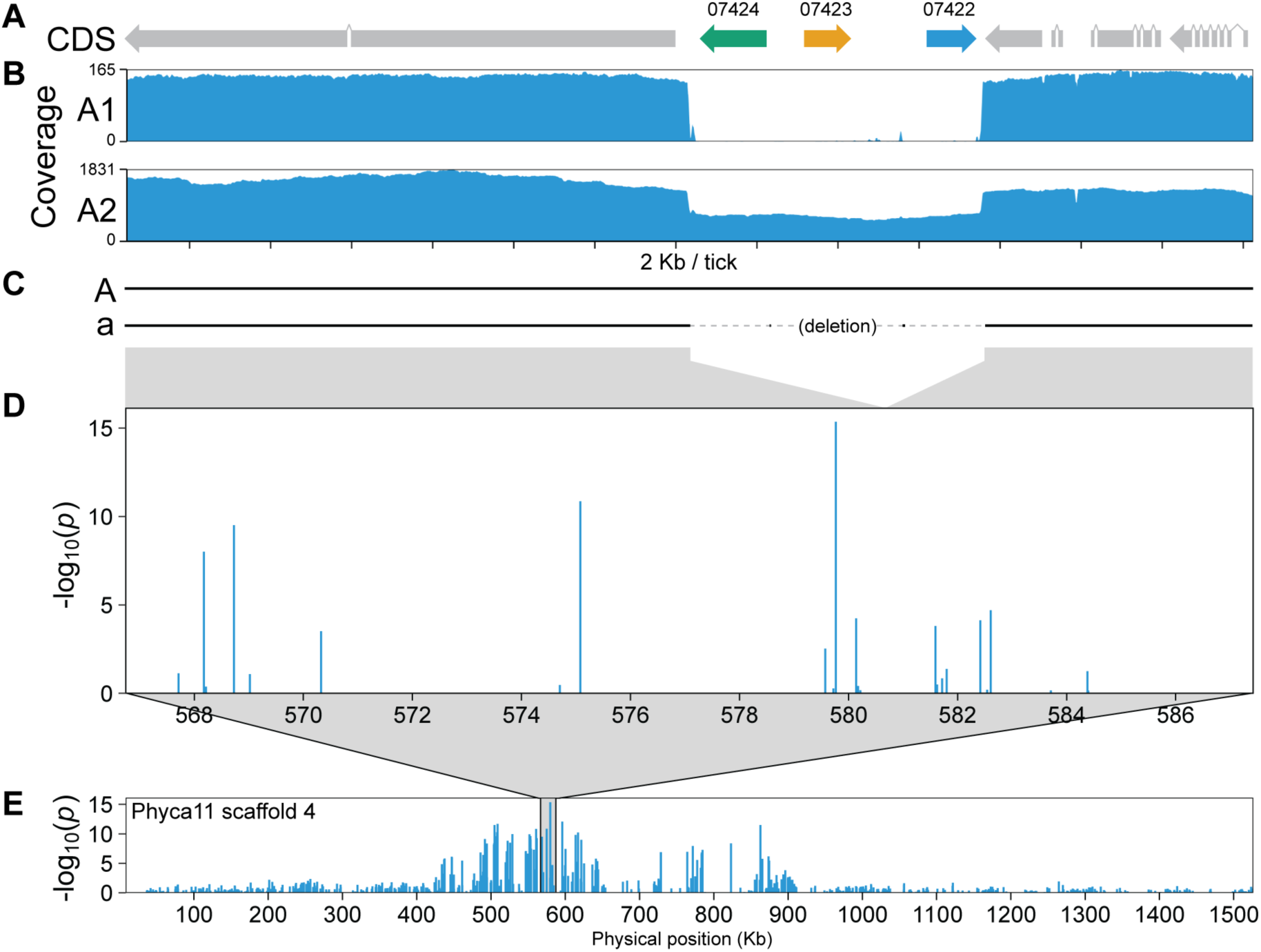
The *Phytophthora capsici* mating-type locus is hemizygous. (A) Gene models annotated on the A haplotype; the mating-type–associated gene cluster is highlighted in color, with genes named by *P. infestans* orthology. (B) Long-read depth across the locus in representative A1 and A2 isolates (55330 and LT1534, respectively) showing homozygous *aa* in A1 (coverage drop to 0 across the A-specific interval) and hemizygous *Aa* in A2 (coverage reduced to ∼50% across the interval), consistent with allele-specific presence/absence. (C) Pairwise alignment of the corresponding interval between the haplotypes of locally phased ASM3032425v1 (A2 strain) assembly. Gray connectors link aligned segments to their physical positions in the association plots. The two assemblies represent alternate mating-type haplotypes and are labeled a and A. (D) Zoomed view of the ∼19-kb interval surrounding the top-associated SNP, showing the same association signal at higher resolution. The Phyca11 genome used for GWAS is haplotype collapsed and corresponds to the hemizygous allele. (E) Genome-wide association signal across scaffold 4 of the Phyca11 assembly, generated using the same genotype set and GWAS workflow as Vogel *et al.,* 2020 (*12*).

The location of the A2-specific cluster is in agreement with raw data from a prior GWAS study focused on mating type that employed a reduced representation sequencing method with the Phyca11 reference genome (Fig. 10D,E) (*12*, *16*). That study supported the hypothesis that A1 strains were homozygous *(aa)* and A2 strains heterozygous *(Aa)* at a mating type locus, but could not define the specific mating determinant. When we reanalyzed that data using a newer assembly, mating type mapped to the three-gene cluster. This cluster was apparently missed in the original GWAS study since the scaffold containing the GWAS signal (scaffold 4) is haplotype-collapsed and bears only the null allele of the locus. The other allele (containing orthologs of PITG_07424 and PITG_07423) was partly assembled as “scaffold 352” but lacked coverage to allow SNP detection due to reduced representation sequencing strategy. Our searches failed to detect similar hemizygosity in other species of *Phytophthora*, but this may need to be investigated again once a greater number of high-quality assemblies are developed for the genus. Still, the apparent plasticity in mating type regulation between *P. capsici* and *P. infestans* is remarkable.

## Discussion

We have identified a three-gene cluster that specifies the A2 mating type in *Phytophthora* by encoding a key enzyme in α2 mating hormone biosynthesis and two proteins with previously unreported domain architecture. Although the α1/α2 signaling system is conserved within the genus (*6*), its functional implementation differs strikingly between species. In *P. infestans*, mating type is regulated by a locus genetically unlinked to the α2 biosynthetic gene, whereas in *P. capsici*, a physical deletion of the cluster determines mating type. The recurrent degradation of this cluster in homothallic species further indicates that homothallism evolved multiple times from heterothallics. Our findings thus reveal unexpected flexibility in how mating type is determined in the genus.

Our functional and metabolic analyses show that α2 is produced from phytol in two steps. The first is performed by PITG_21524, a CYP present in both mating types. The second step is performed by another CYP, PITG_07424. The two are closely related and we propose that the latter arose from a duplication of PITG_21524 which became specialized for α2 production. The two-step pathway is consistent with previously proposed biosynthetic intermediates (*10*).

Further, we show that α2 is only produced by A2 strains because A1s either lack or do not express PITG_07424, depending on the species. We confirmed the role of this enzyme by CRISPR-mediated disruption in an A2 background, which resulted in the conversion of the A2 metabolome and phenotype into A1.

However, PITG_07424 alone is insufficient to fully establish A2 identity, as its overexpression in an A1 background yielded a self-fertile strain rather than the A2 phenotype. This suggests that other genes are involved. Consistent with this, our differential expression analysis identified two additional A2-enriched genes, which have a novel START-FYVE domain architecture. Their physical clustering with PITG_07424 across the Peronosporales order is likely related to their co-regulation, and strongly implies a role in mating. The existence of the cluster is reminiscent of the situation in basidiomycetes where mating type loci encode both the mating pheromones and their receptor (*17*). In *Phytophthora*, the START-FYVE proteins might thus function in hormone deployment or perception.

Our unexpected discovery of structural polymorphism as a determinant of mating type in *P. capsici* explains the GWAS signal in the prior study (*12*) and the report that mating type switching can only occur in A2 to A1 direction (*18*). Our findings are corroborated by a study (*19*) showing that an allele only from the A2 parent was co-inherited with mating type, although no candidate genes were identified due to large linkage blocks. In contrast, in *P. infestans*, and likely other *Phytophthora* species, the fully functional cluster is present in both mating types but is expressed only in A2. This finding means that although the mating-type system originated once (*6*), the underlying genetic determinants have diverged during the radiation of the Peronosporales, exemplifying developmental system drift and regulatory plasticity of the mating system (*20*, *21*).

In contrast to the heterothallics, over a half of homothallic (self-fertile) species in *Phytophthora* bear disruptions of at least one gene in the cluster and those events are not uniform. This is evidence that homothallism evolved from heterothallism on multiple occasions, and is consistent with the observation that homothallic species are interspersed among heterothallics in molecular phylogenies (*22*). In some instances, self-fertility may have evolved through changes outside of the cluster that enable the sexual program to be activated, rendering the cluster dispensable and thus prone to degeneration. In the case of the homothallics that maintain an intact three-gene cluster, the change may have occurred such that those genes are still required. However, homothallism in *Phytophthora* does not seem to involve phenomena observed in fungi such as the acquisition of multiple mating type loci (as in homothallic *Aspergillus* species (*23*)) or genome arrangements that affect their expression as in the case of *Saccharomyces* (*24*). Since how the hormones stimulate mating in *Phytophthora* remains to be determined, it is possible that mutations outside the cluster have broken a self-incompatibility barrier, resembling how plants such as *Arabidopsis thaliana* evolved self-fertility by disruption of self-incompatibility genes (*25*).

While our results independently validate α1/α2 hormones as the main triggers of oospore formation in heterothallic *Phytophthora*, they also point towards additional hormonal complexity. Notably, the A1:F3 derivative of phytol, which has not been previously described, attracts the growth of A2 strains but without inducing oospores. This resembles a report that synthetic α1 also induces directional growth along with oospore formation in A2 cultures (*26*). Because A1 strains cannot produce α1 until exposed to α2, it is possible that A1 strains initially release A1:F3 to recruit A2 strains. This would be analogous to mating chemotropism in yeast (*27*). The requirement for key A1-derived metabolites and possibly gradients or other contextual cues may help explain why the responses of A2 strains to exogenous α1 are often inconsistent (*26*).

Overall, our findings prompt reconsideration of how mating is regulated across species boundaries. The demonstration of mating hormone universality within *Phytophthora* led to an expectation that the genetic determinants of mating type would also be universal (*5*, *6*, *28*), but the contrast between species like *P. infestans* and *P. capsici* shows this to be untrue. The next step in understanding the sexual cycle would be identifying the receptor(s) of the mating hormones and their downstream signaling cascades. The latter may also exhibit divergence within *Phytophthora* since the nature of gametangial interactions varies between species. Our findings also offer a path toward practical tools. Disease management relies on monitoring populations of the pathogen to assess the likelihood of genetic variation and whether durable over-wintering oospores may form within a field. A molecular test for mating type based on the genes described here would replace the laborious assays that are now used, which require tester strains and the purification of strains from field material.

## Acknowledgments

We thank Audrey Ah Fong for providing valuable advice and guidance. This research was supported by an award to HSJ from the National Institute of Food and Agriculture of the United States Dept. of Agriculture, and the Singletary Endowed Chair in Agriculture.

## Acknowledgments

We thank Audrey Ah Fong for providing valuable advice and guidance.

## Supplementary Materials

### Materials and methods

#### Phytophthora strains and culture conditions

*P. infestans* strains 1306 (A1 mating type) and 618 (A2 mating type), and *P. nicotianae* strains 1751 (A1) and 3881 (A2) were obtained from the UCR *Phytophthora* collection. Cultures were maintained on rye B media with 1.5% agar, supplemented with 50 μg ml^-1^ penicillin G and 10,000 units ml^-1^ nystatin (*29*). Liquid cultures were grown in rye B broth clarified by centrifugation at 10,000 × g for 15 min, containing the same antimicrobials. Mating type and hormone assays were performed on clarified rye with 1.5% agar.

To identify phytol metabolites in an untargeted manner, A1 and A2 *P. infestans* strains were grown in broth supplemented either with 1 ml l^-1^ ethanol (control) or with 1 ml l^-1^ ethanol containing phytol (≥97% Sigma-Aldrich, Cat. No. W502200) at 1 mg ml^-1^, yielding a final concentration of 3.4 mM. For each strain and treatment, five independent cultures were established, each consisting of 80 ml of broth per 150 mm plate. Plates were inoculated with sporangia at a concentration of 1.5 × 10^5^ per ml, sealed with plastic film, and incubated for 5 days at 19°C in the dark. Cultures were harvested by vacuum filtration followed by a water rinse, and flash-frozen in liquid nitrogen. Metabolites were immediately extracted as described below.

Histone deacetylase assays were conducted in clarified rye broth with five replicates using ethanol stock solutions of phytol alone or phytol combined with trichostatin A (Thermo Scientific, Cat. No. AC471120050), each applied to a final concentration of 1 mg l^-1^. RNA was then extracted from mycelia and media were used for statistical metabolomic analysis.

To facilitate identification of phytol-derived metabolites, a 100 ml culture of the two strains was prepared containing either unlabeled or deuterium-labeled phytol (Toronto Research Chemicals Inc, Cat. No. TRC-T398919) at 3.4 mM. Phytol-supplemented broth cultures were used for metabolic profiling of knockout, overexpressing, and wild-type strains were established in parallel with wild-type A1 and A2 strains. For metabolite purification, larger-scale (2.3 l) cultures of each strain (18 plates, 150 mm diameter) were used.

#### CRISPR/Cas12a knockouts

Knockout mutants of PITG_07424 were generated in the A2 *P. infestans* strain 618 using our previously established CRISPR/Cas12a system (*30*). Briefly, four guide RNAs targeting sequences near the 5′ end of the PITG_07424 coding sequence were designed, synthesized, and cloned into a CRISPR/Cas12a expression plasmid carrying a G-418 resistance marker. Protoplast preparation, transformation, and regeneration were performed as described (*30*). Following transformation, protoplasts were plated on regeneration medium supplemented with G-418 at 8 µg ml⁻¹, and colonies emerging under selection were isolated. Putative transformants were screened by PCR using primers listed in Table S4, and PCR products spanning the target region were Sanger-sequenced to confirm disruption of PITG_07424. Verified knockout mutants were subsequently grown on medium lacking selection.

#### CYP cloning and expression

*P. infestans* CYP genes were amplified from cDNA or genomic DNA, using primers (Table S4) designed for restriction enzyme-based cloning or assembly using the NEB HiFi DNA assembly system (New England Biolabs, Ipswich, MA, USA). For expression in *S. cerevisiae,* PCR products were cloned into pESC-T or pESC-L with or without a C-terminal FLAG using *E. coli* DH5α cells. After verifying the constructs by whole-plasmid sequencing, yeast *S. cerevisiae* WAT11 strain was transformed as described (*31*). For co-expression of two genes, yeast was doubly transformed with the desired combination of constructs and selected on the appropriate dropout medium. WAT11 expresses the *Arabidopsis thaliana* CPR gene to enhance P450 catalytic efficiency (*32*). Transformants were grown in 5 ml of dextrose dropout medium to 3x10^7^ cells ml^-1^, washed twice with 5 ml of water, and resuspended in 10 ml of galactose dropout medium to induce GAL1/10 promoter in the pESC plasmids. After vortexing, the culture was split into two 5 ml cultures to which 5 µl of ethanol, or 1 mg ml^-1^ phytol in ethanol, was added. After 72 h, cells and media were collected by centrifugation. Cells were washed with 5 ml of water before metabolite extraction. CYP production was confirmed by extracting proteins from a sample of the cell pellet (*33*) followed by western blotting with anti-FLAG monoclonal antibody (Sigma-Aldrich, Cat. No. F1804) as described (*34*).

The same restriction enzyme cloning strategy was used to generate ectopic expression constructs for *P. infestans*, using the pTOR vector with G-418 selection. For constitutive overexpression, CYP coding sequences were placed under control of the *Ham34* promoter within pTOR; for constructs driven by the native promoter, the *Ham34* promoter was excised from pTOR by double digest with *Kpn*I/*Not*I restriction enzymes, and the native promoter-CDS fragment was ligated in its place. Transformation was performed as described above.

#### Metabolite extraction and LC-MS

Liquid culture media of *Phytophthora* and yeast were extracted two times with an equal volume of ethyl acetate. The pooled organic phases were washed with 0.1 volume of water, dried under a gentle stream of nitrogen, and the residues resuspended in acetonitrile containing 0.1% (v/v) formic acid. After centrifuged at 17,000 × g for 10 min, the supernatant analyzed by LC-MS. Mycelia was harvested by vacuum filtration through Whatman 54 filter paper, and ground to a fine powder under liquid nitrogen. The ground tissue was extracted with methanol by vortexing and 15 min of sonication in an ultrasonic bath (Branson 5800). Extracts were centrifuged at 17,000 × *g* for 10 min, and the supernatants used for LC-MS analysis. Yeast cell pellets were resuspended in 200 µl methanol and mixed with an equal volume of 0.5 mm glass beads. Cells were disrupted using a bead beater (2010 Geno/Grinder, SPEX SamplePrep LLC) with two 2 min cycles at 20 Hz, and 30 s rest time. Lysates were centrifuged at 17,000 × g for 10 min, and the supernatants were collected for LC-MS analysis.

Metabolite profiling was performed using a Vanquish UHPLC system (Thermo Scientific) with a ZORBAX RRHD Eclipse Plus C18 Column (Part Number: 959757-902, Agilent) coupled to an Orbitrap IQ-X mass spectrometer equipped with a heated electrospray ionization source. Chromatographic separation was carried out at 35°C. The autosampler was maintained at 10°C with a draw and dispense speed of 5 µl s⁻¹, using a pre- and post-injection needle wash (2 s wash time at 24 µl s⁻¹). Mobile phase A consisted of 5% acetonitrile with 0.1% formic acid, and mobile phase B was acetonitrile with 0.1% formic acid. The flow rate was 0.30 ml min⁻¹. The chromatographic gradient was as follows: the run began at 30% B. The proportion of B was increased linearly to 100% at 14 min and maintained at 100% from 17 min. Between 17 and 18 min the system was returned to 30% B to re-equilibrate the column. The total runtime was 20 min. A diode-array detector simultaneously recorded UV spectra from 190 to 800 nm at 10 Hz. Mass spectrometric detection was performed in positive-ion mode. The spray voltage was set to +3500 V, sheath, auxiliary, and sweep gases were set to 50, 10, and 2 arbitrary units, respectively, the ion transfer tube was maintained at 275°C, and the vaporizer temperature was 350°C. Internal calibration using EASY-IC was enabled throughout the run. Full MS scans (MS¹) were acquired in the Orbitrap at a resolution of 60,000 (m/z 200) over a scan range of m/z 115–1500. The AGC target was 4 × 10⁵, the maximum injection time was controlled automatically, one microscan was collected per scan, and data were recorded in profile mode. Data-dependent MS² scans were triggered for precursor ions exceeding the dynamic intensity threshold (minimum intensity 25,000; relative threshold 20%). MS² spectra were acquired in the Orbitrap at a resolution of 30,000 using quadrupole isolation with a 1.6 m/z window and HCD fragmentation with stepped collision energies of 15%, 40%, and 55%. The AGC target was 5 × 10⁴ and injection time was set automatically (≤54 ms). Three MS² scans were collected per cycle. MS³ scans were subsequently triggered from MS² fragments using a 2.5 m/z isolation window and HCD with a fixed collision energy of 30%. MS³ spectra were also acquired at 30,000 resolution with an AGC target of 5 × 10⁴ and automatic injection control. Both MS² and MS³ spectra were collected in centroid mode.

Raw files were converted to centroid mode using MSConvert (ProteoWizard) with default peak-picking settings appropriate for Orbitrap data. Processed files were subsequently imported into MZmine for mass feature detection, deconvolution, isotope recognition, and chromatogram building using the program’s default parameters. The resulting feature table, containing mass features, retention times, and peak areas, was exported and uploaded to MetaboAnalyst for statistical analysis. Within MetaboAnalyst, data were normalized using quantile normalization, log₂-transformed to stabilize variance, and scaled using autoscaling (mean-centering followed by division by standard deviation). This enabled downstream identification of statistically significant and differentially accumulated metabolites. To assign putative molecular formulas and predict the chemical class of metabolites, spectral data were analyzed using SIRIUS. Molecular formula prediction was performed by combining isotope pattern scoring, MS¹ accuracy, and fragmentation tree computation in CSI:FingerID. Structure prediction and metabolite class assignment were generated using CANOPUS, which applies machine-learning models trained on large spectral libraries to classify compounds within the ChemOnt ontology.

Fragmentation trees were constructed from MS² and MS³ spectra to refine formula selection and improve annotation confidence for unknown features. Since the sodium adduct ion did not fragment in MS^2^, related ions were identified including a protonated ion matching the dissociation of one or two hydroxyls. Fragmentation was therefore performed on the dehydrated ions [M-H_2_O+H]+ or [M-2H_2_O+H]+. Raw data files were inspected as needed in FreeStyle 1.8 SP2 QF1.

To help confirm the identity of metabolites produced by cytochrome P450 enzymes heterologously expressed in yeast, their retention times were compared with those extracted from Phytophthora cultures run sequentially on a well-equilibrated LC-MS system. The extracted ion chromatograms were also compared and structural identity evaluated by mass spectrometry, including MS^2^ and MS^3^ fragmentation analyses. Also, yeast-derived samples were spiked with Phytophthora-derived metabolites and checked for co-elution.

#### Analysis of phytol metabolites

To determine the regiochemistry of cytochrome P450-mediated oxidations of phytol by *Phytophthora* enzymes, isotopically labeled phytol containing six deuterium atoms (phytol-*d*6) was used as a substrate. Deuterium atoms were positioned at C2, C4, and C20, enabling discrimination between oxidative modifications occurring at the α-end of the phytol molecule versus the distal ω-end. This further allowed assignment of phytol-derived metabolites detected in A1 and A2 strains. Feeding experiments with phytol-*d*6 were performed under identical conditions as those used for unlabeled phytol. Following incubation, metabolites were extracted as described previously and analyzed by LC-MS using the same chromatographic and mass spectrometric parameters. Incorporation and positional retention of deuterium were further validated by molecular formula prediction based on high-resolution MS spectra acquired with a resolving power of 500,000 at m/z 200.

Girard’s T reagent (Thermo Scientific Chemicals, AC119870500) was used to derivatize carbonyl-containing metabolites in *Phytophthora* extracts to assess the presence of keto functional groups in phytol-derived metabolites. Girard’s T reagent selectively reacts with ketones to form permanently charged hydrazone derivatives. For derivatization, 250 µl of metabolite extract was diluted with 750 µl of HPLC-grade water, followed by addition of 50 µl acetic acid. After gentle mixing, the solution was transferred to tubes containing 50 mg of Girard’s T reagent and incubated at 85°C for 4 h. Reactions were quenched by addition of 220 µl of 4 M NaOH and diluted with 2 ml HPLC-grade water to a volume of approximately 3 ml.

Solid-phase extraction (SPE) was then performed using Strata-X cartridges (Supelco), which were conditioned with 1 ml methanol and equilibrated with 1 ml HPLC-grade water. Samples were loaded, washed with 1 ml 5% methanol in water, and eluted with 1 ml 2% formic acid in methanol. Eluates were dried and reconstituted for LC-MS analysis.

#### Metabolite fractionation

Three most abundant phytol-derived compounds from each mating type (six fractions in total) were selected for fractionation (Fig. 4) from the larger-scale culture media extracts. Chromatographic separation of A2 metabolites was carried out on a XBridge BEH C18 Column, 130Å, 3 mm × 100 mm, 3.5 µm column (Cat No. 186003027, Waters), at a flow rate of 0.90 ml min⁻¹ using solvent A as water +0.1% FA and solvent B as acetonitrile + 0.1% FA. The gradient began at 38% B at 0 min and transitioned to 55% B at 12 min. The mobile phase was increased to 100% B at 13 min and held until 15 min before returning to initial conditions (38% B) at 15.5 min and holding until 20 min for re-equilibration. Chromatographic separation of A1 metabolites was carried out on a InfinityLab Poroshell 120 EC-C18, 3.0 mm × 50 mm, 2.7 μm column (Cat. No. 699975-302T, Agilent) at a flow rate of 0.30 mL min⁻¹ using solvent A as water +0.1% FA and solvent B as acetonitrile + 0.1% FA. The gradient began at 33% B at 0 min and transitioned to 85% B at 10 min. The mobile phase was increased to 100% B at 11 min and held until 13 min before returning to initial conditions (33% B) at 13.5 min and holding until 15 min for re-equilibration. In both cases, a VWD detector monitored absorbance at 200 nm with a peak width setting of ≥0.1 min. Metabolite profiling on the Agilent 6224 TOF LC/MS system was performed using a Dual ESI source operated in positive-ion mode with centroid data acquisition. The drying gas temperature was maintained at 325°C at a flow rate of 10 l min⁻¹, and the nebulizer pressure was set to 30 psi. The auxiliary gas was held at 300°C with a flow of 3 l min⁻¹. The capillary voltage (Vcap) was 3500 V. The fragmentor voltage was set to 175 V, with the skimmer at 65 V and the Oct 1 RF at 750 V. MS data were collected with a 5 spectra s⁻¹ acquisition rate using a fixed m/z acquisition range of 100–1000. After establishing the method and confirming the reproducibility of the retention times, the flow was redirected from the mass spectrometer into the waste container or fraction collection tubes. Three fractions were collected from A1 media extract (F1: 6–6.8 min, F2: 7–8 min, F3: 8.2–9.4 min), and three from A2 extract (F1: 6–7 min, F2: 11.5–12.2 min, F3: 12.2–13 min). Fractions were freeze-dried, centrifuged, and resuspended in ethanol. Fraction purity was assayed using the Orbitrap IQ-X LC-MS system as described above and the results are shown in Fig. S17.

#### Hormone bioassays and mating assays

Twenty microliters of each fraction in ethanol (or solvent alone) were pipetted on paper disks supported by a needle. After evaporation of the solvent, the discs were inverted in front of the growing edge of the *Phytophthora* culture. To assess whether purified metabolites were transformed during the assay, the disks were collected after the experiment, extracted by vortexing for 10 min at maximum speed in 1 ml ethyl acetate, and analyzed by LC-MS, using MZmine to detect mass features and quantify peak areas. To focus on phytol-derived metabolites, the feature list was restricted to ions in the m/z 250–400 range and molecular formulas with 20 carbons. The resulting feature table was then manually curated to remove artifacts and redundancies, and peak areas extracted for comparison across samples. Finally, more information on the detected features was obtained by matching the paper-disc features (accurate mass and retention time) to features detected in our *P. infestans* metabolomics datasets and in phytol-*d*6 feeding experiments.

Identity between α2 hormone of *P. infestans* and previously reported α2 hormone in *P. nicotianae* was established via bioassay as previously described (*6*). Application of *P. infestans* α2 to a culture of A1 strain of *P. nicotianae* induced abundant oospore formation, confirming that α2 isolated in the present study is biologically equivalent to the canonical α2 hormone (Fig. S18). No oospores were observed in solvent-only or other-fraction controls.

Mating assays were performed as described (*35*). In brief, 3 mm-wide agar strips were excised from 7-day-old cultures of the strains being tested and placed 1.5 cm apart on a fresh agar plate. After two weeks, oospore production was evaluated by excising a 1-cm-wide agar strip from between the two inoculum strips, which was sectioned and examined under a microscope for oospores. Self-fertility of all strains and mutants was assessed by microscopic examination of two-week-old cultures.

#### Phylogenetic analysis and ortholog identification

A species tree of the Peronosporales was constructed based on concatenated sequences of BUSCO genes, starting from all (n=313) assemblies in NCBI Genome as of the time of analysis (June 2025), 14 *Phytopythium* genomes, and four high quality *Pythium* genomes as an outgroup (*P. oligandrum*, *P. cedri*, *P. myriotylum*, *P. insidiosum*). As most genomes lacked gene models, we used BUSCO v6.0.0 with the eukaryota_odb12 database in genome mode to extract BUSCO genes. We removed genomes which had excessive missing, fragmented. or duplicated BUSCOs. Then, we selected BUSCOs that were missing in <5, fragmented in <6, duplicated in <15 genomes. This resulted in 80 genes (Table S5) and 254 genomes. This was trimmed to 114 species (Table S6) based on those with the most complete single copy BUSCOs, and best N50 values. After aligning each BUSCO using MAFFT (v7.505) in G-INS-i mode and removing poorly aligned or duplicate sequences, the alignments were trimmed using clipkit (--mode kpic-gappy --gaps 0.2), and concatenated and analyzed using IQ-TREE with each BUSCO gene inputted as a partition. Extended ModelFinder with automatic merging of partitions was used to select the best-fit substitution model for each partition and merge partitions with similar model parameters. Branch support was assessed using 10,000 bootstrap replicates. To reduce systematic error from compositional heterogeneity, we enabled the IQ-TREE symmetry/compositional test and removed partitions failing this test.

This tree was then used as a species-tree in an ortholog mining pipeline to identify CYP genes from the 114 genomes. This used tblastn with an *E*-value cutoff of 1e-5) against oomycete CYP proteins reported previously (*14*). Hits with <30% identity and lengths <200 amino acids were removed. Genomic regions with CYP hits, including ±200 nt flanking sequence, were extracted, aligned with MAFFT in L-INS-i mode, and used to annotate the start codon. Open reading frames were extended *in silico* to the first in-frame stop codon, translated, and truncated protein sequences (length <80% of the alignment) removed. One CYP gene known to contain an intron was manually curated to correctly reconstruct the full-length coding sequence. In rare cases, where 200 nt flank was not enough to cover the full CDS, it was extended further. The resulting CYP protein set, together with the genome-wide species tree, was used as input to OrthoFinder v3.1.0 for gene-tree/species-tree reconciliation and inference of orthogroups and duplication events. The final reconciled gene tree is shown in Fig. S2. .

Orthologs (or their remnants) of *P. infestans* PITG_07422 and PITG_07423 in the 114 genomes were identified by reciprocal BLAST using an *E*-value cutoff of 1e-5, followed by the reciprocal best hit method. Apparent false positives based on low identity or coverage were removed. Apparently truncated hits were then extended as described for the CYP orthologs. In genomes with no hits, a second search was performed using protein sequences from the most closely related species which had an ortholog followed by reciprocal blast. To check for synteny, genomic regions flanking the annotated features were extracted using a custom Python script. Annotated features within 10 Kb were merged, and the resulting sequences and coordinate-adjusted GFF3 annotations used for data visualization (Fig. S2). Synteny was identified by visual inspection.

#### Expression analysis

RNA-seq reads (Table S7) from eleven A1 mating-type strains (1306, 8811, 88069, Sw1, Sw3, F80029, Pink6, M4, M60, M66, M108) and ten A2 mating-type strains (618, E13, Sw2, Sw4, IPO82001, 3928A, 4432, M70, M76, M738), were adapter- and quality-trimmed with BBDuk (BBMap v39.01) and aligned to the *P. infestans* genome using STAR (v2.7.11b, default settings, max intron length set to 5 Kb). The genome used was a telomere-to-telomere assembly of *P. infestans* strain 1306, which we annotated by lifting over gene models from the T30-4 assembly using liftoff (v1.6.3, default settings). As some of the data came from paired-end sequencing, only their first read was used. Gene-level counts were obtained with featureCounts (Subread R package, v2.1.1). Principal component analysis (PCA) was used to assess data quality. Differential expression between A1 and A2 strains was assessed using edgeR (v4.8.1), with library-size normalization and a negative binomial generalized linear model incorporating the sequencing study as an independent variable. Counts of replicates were averaged before the analysis. Multiple testing was controlled using a edgeR false discovery rate (FDR) as with a cutoff threshold of ≤0.05. Strength of the association was further evaluated by examining normalized CPM values across the A1 and A2 strains. Genes of interest were also tested by RT-qPCR (*36*) using the shown in Table S4.

To enable detection of differential transcription from unannotated or misannotated regions, we also performed an annotation-free analysis using DERfinder (v1.45.0), following the default workflow. Briefly, genome-wide per-base coverage was computed from STAR-aligned BAM files, expressed regions were defined and summarized per sample, and differential expression between A1 and A2 strains was tested by DESeq2 in a model including sequencing study as a covariate with FDR control (≤0.05).

#### Functional annotation of genes

Functional annotation of genes was performed using the NCBI Conserved Domain Search tool, combined with transmembrane topology analysis (Phobius, TMHMM), localization prediction (DeepLoc 2.1) and AlphaFold 3. 3D alignment of AlphaFold 3 models with each other or with crystallography-derived 3D models from PDB (START and FYVE domain PDB accessions 1em2 and 1vfy, respectively) was performed using RCSB protein Data Bank Pairwise Structure Alignment webserver (jFATCAT rigid). Protein domain architectures containing both START (PF01852) and FYVE (PF01852) domains, in any order, were identified using the InterPro webserver.

#### Promoter analysis

For phylogenetic footprinting of the CYP promoter, we extracted from all species with an intact three-gene cluster the intergenic region between its ortholog and adjacent START-FYVE genes (PITG_07423 in *P. infestans*). This corresponding to ∼1 kb upstream of the CYP start codon. These were aligned using Pro-Coffee with default parameters, and the resulting multiple sequence alignment was used to identify conserved elements. To complement the alignment-based approach, we used MEME to search for conserved motifs with zero or one occurrences per sequence across all orthologous promoters. For intraspecific promoter comparisons in *P. infestans*, we used phased genome assemblies of two A1 strains (1306 and 6629) and two A2 strains (618 and 550), yielding eight promoter haplotypes which were aligned using MAFFT L-INS-i with default settings. Allele-specific expression of the genes of interest was assessed by visual inspection of RNA-seq read mappings of the same strains and their progeny in JBrowse2. The F1 progeny, along with the corresponding DNA and RNA sequencing data used in this study is listed in the Table S7 and have been described previously (*36*). DNA and RNA reads were mapped to the *P. infestans* reference genome as described above, and informative DNA SNPs were identified in RNA reads to confirm expression of both alleles.

#### Reanalysis of P. capsici GWAS

Mapping of available short and long DNA reads (Table S7) of *P. capsici* A1 strains (BYA5, 55330) and A2 strains (LT1534, LT263) was performed against the chromosome level assembly (ASM3032425v1) of an A2 strain LT263. PacBio HiFi, PacBio and Oxford nanopore reads were mapped using minimap2 (v2.30) using map-hifi, map-pb, and map-ont presets, respectively. Short reads were quality- and adapter-trimmed using BBDuk (BBMap v39.01) and mapped using bwa-mem2 (v2.2.1). Similarly, RNAseq reads for strains BYA5, LT263 and LT1534 (Table S7) were trimmed and mapped to the same genome, using the same method as for DGE analysis. Read alignments were inspected in JBrowse2 at the previously identified 3-gene cluster. The genomic region harboring the mating type GWAS signal was localized by mapping the corresponding interval from the original assembly used for GWAS (Phyca11) onto the chromosome-level reference using minimap2 (asm20 preset). The cluster presence was also checked in genomes without publicly available raw reads (Table S6) using blastn.

---

**Fig. S1.**
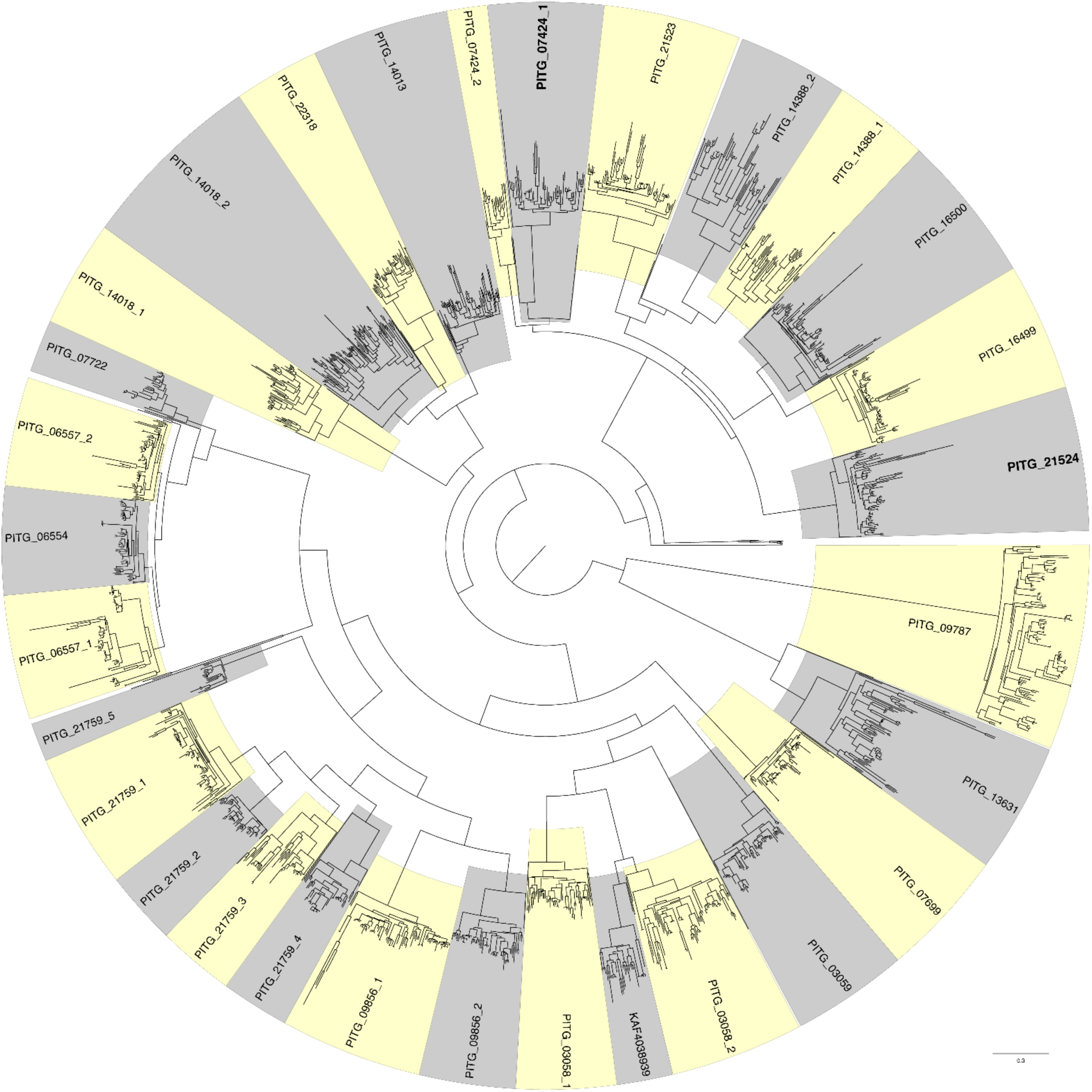
Circular phylogram of all detected Peronosporales cytochrome P450 (CYP) orthologs. Clades representing othogroups are highlighted in alternating yellow and gray colors and are labeled based on the closest gene present in *P. infestans*. An additional number was added as suffix when multiple orthogroups have same closest gene in *P. infestans*. Orthogroups containing CYPs involved in α2 biosynthesis are labeled in bold.

**Fig. S2.**
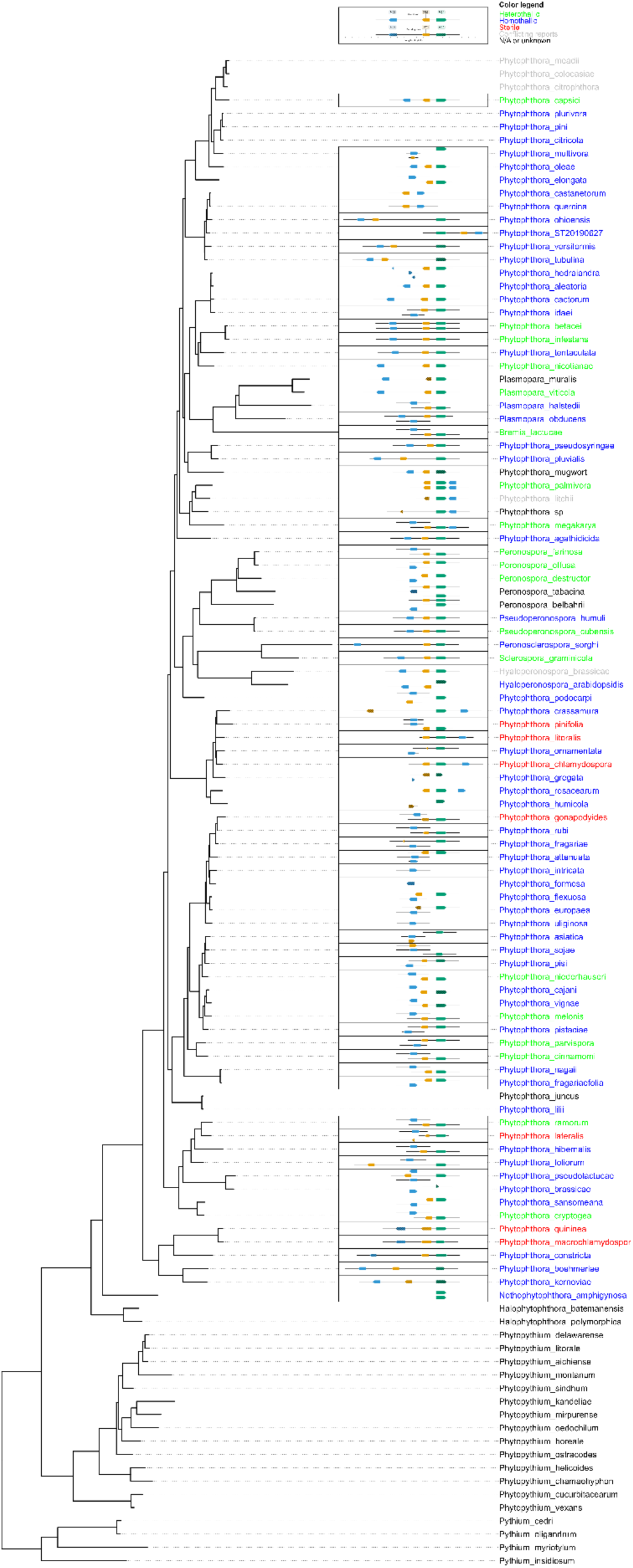
BUSCO-derived species phylogenetic tree with 07424, 07423, and 07422 positions shown for each species where the orthologs were detected. For each gene shown, a 2 Kb flanking sequence is drawn, unless the contig end is closer. Colors of species names indicate the mating system.

**Fig. S3.**
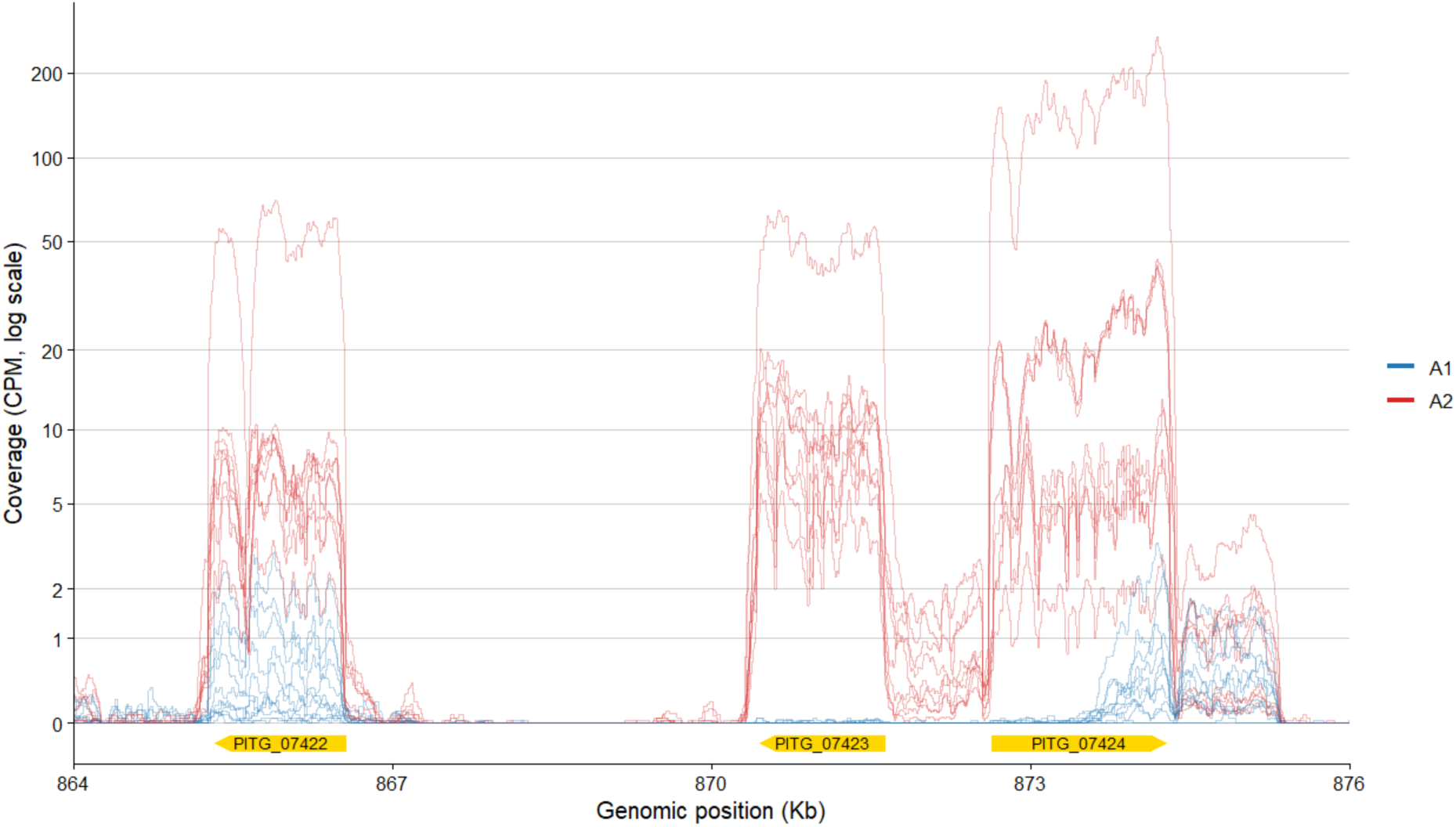
RNA-seq coverage of the gene cluster region for all A1 (blue) and A2 (red) strains used in differential gene expression analysis, with gene annotation at the bottom. Horizontal axis shows genomic position (Kb) on chromosome 13, while vertical axis shows DESeq2 normalized coverage, at logarithmic scale.

**Fig. S4.**
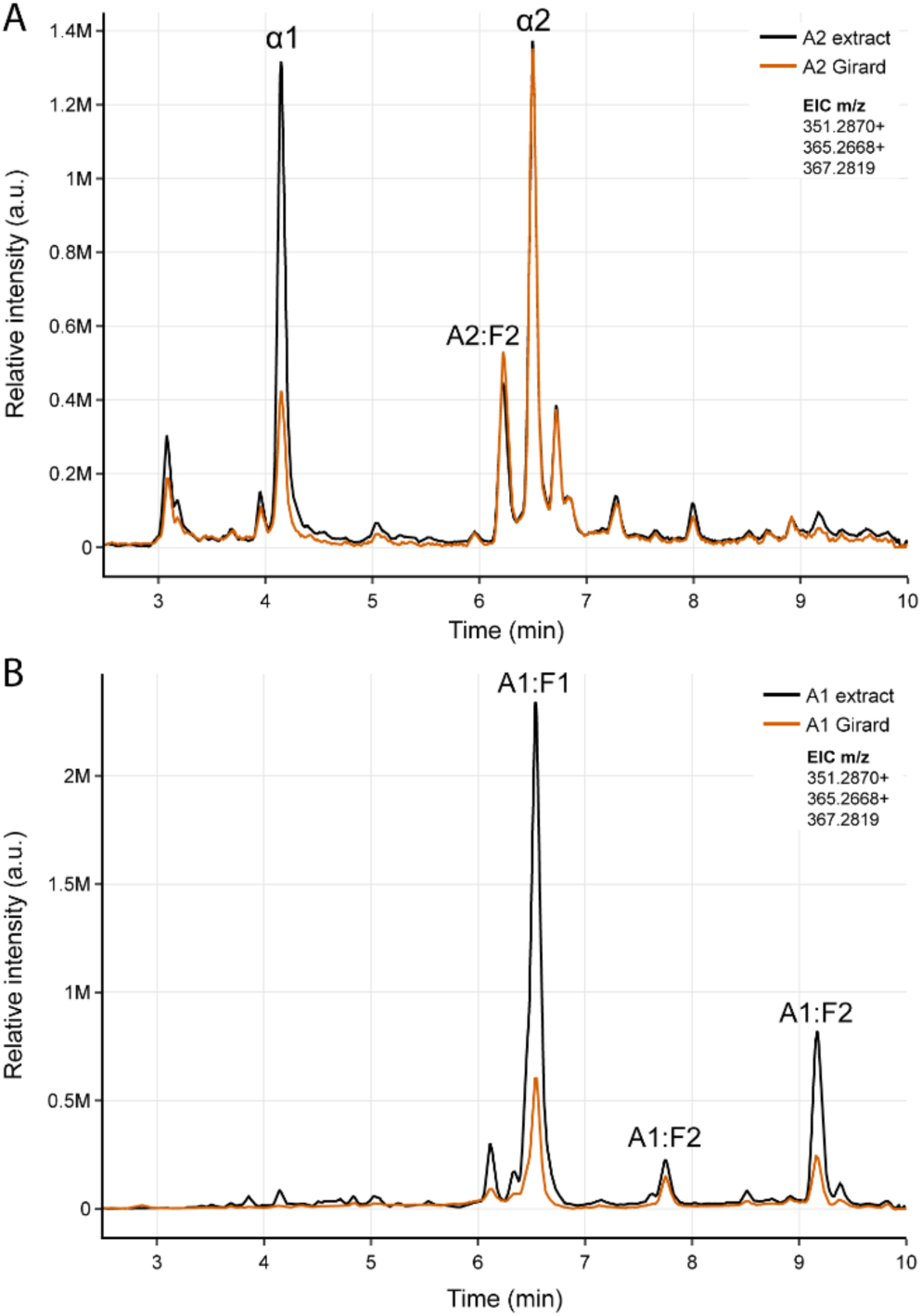
Extracted ion chromatograms (EICs) of phytol-derived metabolites in (A) A2 and (B) A1 media extracts before and after Girard’s T derivatization. Overlaid traces show the untreated extract (orange) and the Girard’s T–treated extract (black). EICs correspond to m/z 351.2870, 365.2668, and 367.2819. Peaks corresponding to fractioned metabolites are indicated above the peak. Reduction of the peak intensity indicates reaction with Girard’s T reagent, consistent with the presence of a carbonyl (keto) functional group in that metabolite.

**Fig. S5.**
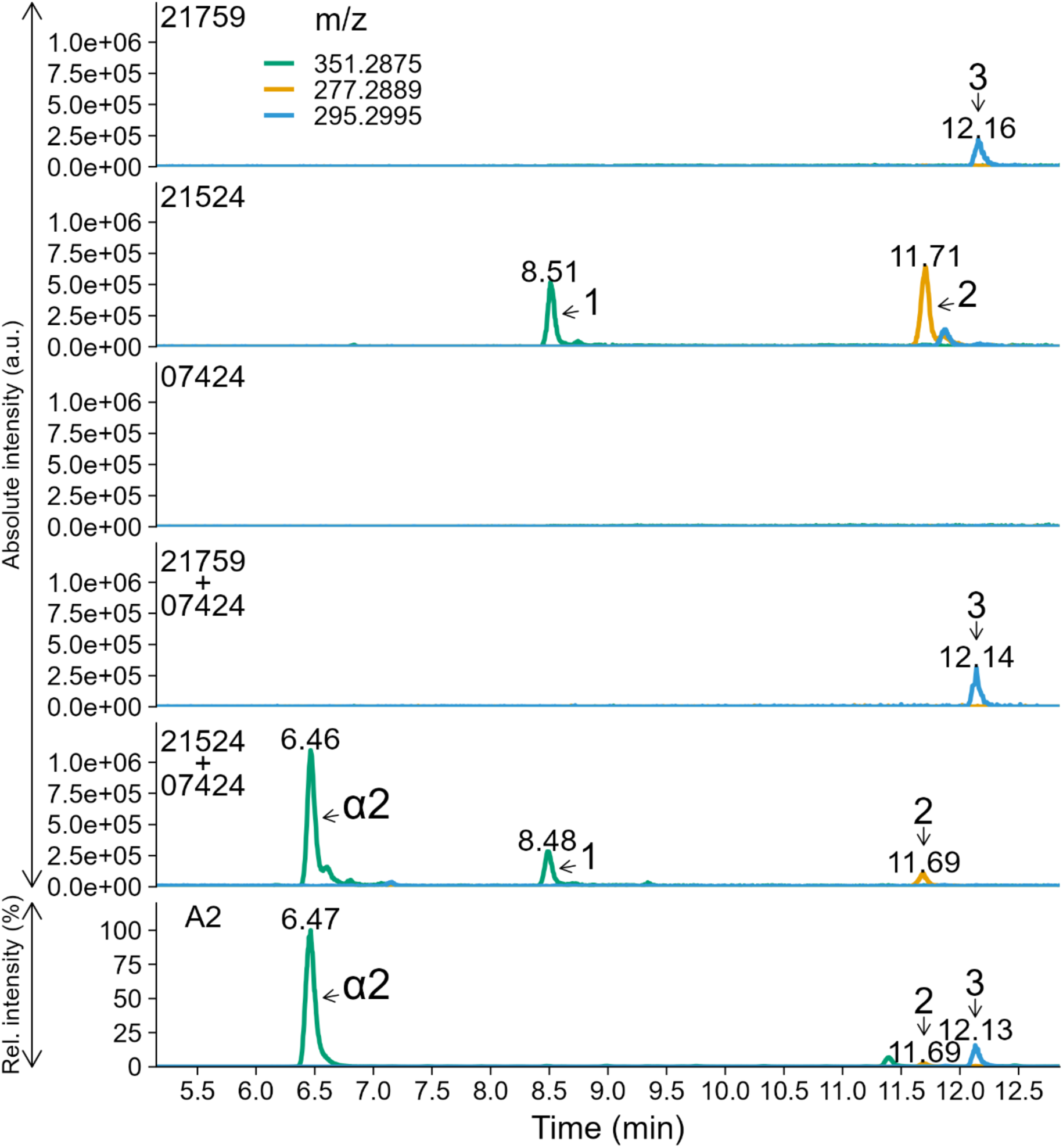
Extracted ion chromatograph (EIC) of media extract from *Saccharomyces cerevisiae* expressing *Phytophthora infestans* cytochrome P450 (CYP) genes 21524, 21579, 07424, or combination (21524+07424 and 21579+07424), showing production of phytol-derived metabolites. For comparison, the *P. infestans* A2 culture EIC is shown below. Phytol was added to all cultures. Traces are shown for m/z 351.2875 (1; green; putative dihydroxyphytol), 277.2889 (2; orange; putative hydroxyphytol), and 295.2995 (3; blue; putative hidroxyphytol). Retention times (min) are indicated above peaks and arrows label peak identities. Yeast EIC panels use identical absolute intensity scaling; the *P. infestans* A2 EIC is displayed on a relative intensity scale.

**Fig. S6.**
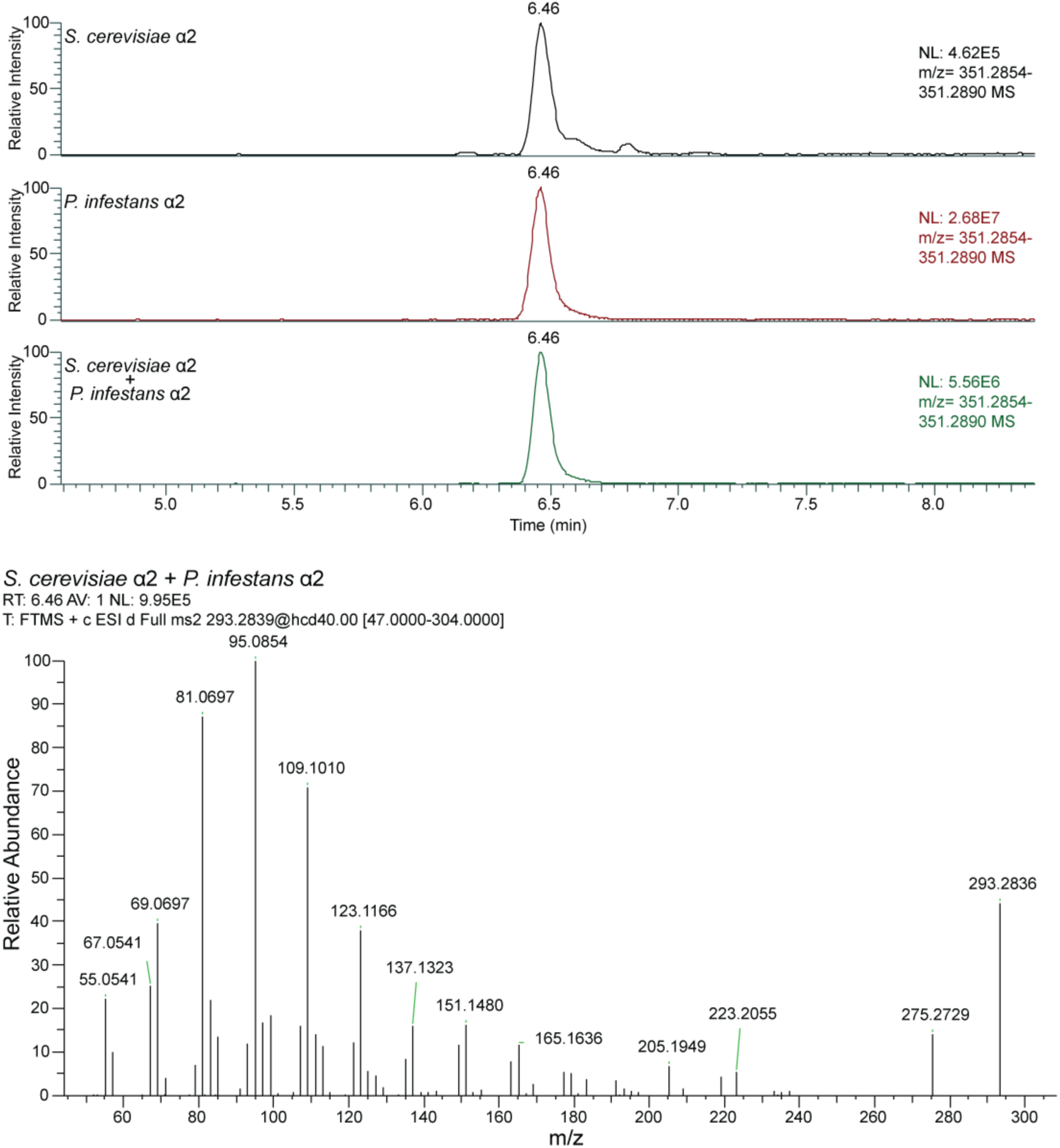
Chromatographic co-elution and MS^2^ validation of α2 metabolites from S. cerevisiae and P. infestans. Extracted ion chromatograms (EICs; m/z 351.2854–351.2890) of α2 from *S. cerevisiae* (top), *P. infestans* (middle), and a co-injected mixture (bottom) are shown. The metabolite elutes at 6.46 min in all samples, demonstrating identical retention times under identical chromatographic conditions. The MS^2^ spectrum of the co-injected sample (bottom panel) shows a single fragmentation pattern corresponding to the precursor ion (m/z 293.2839, HCD 40), matching the fragmentation profile observed for the individual samples (shown in the main text figure 3C).

**Fig. S7.**
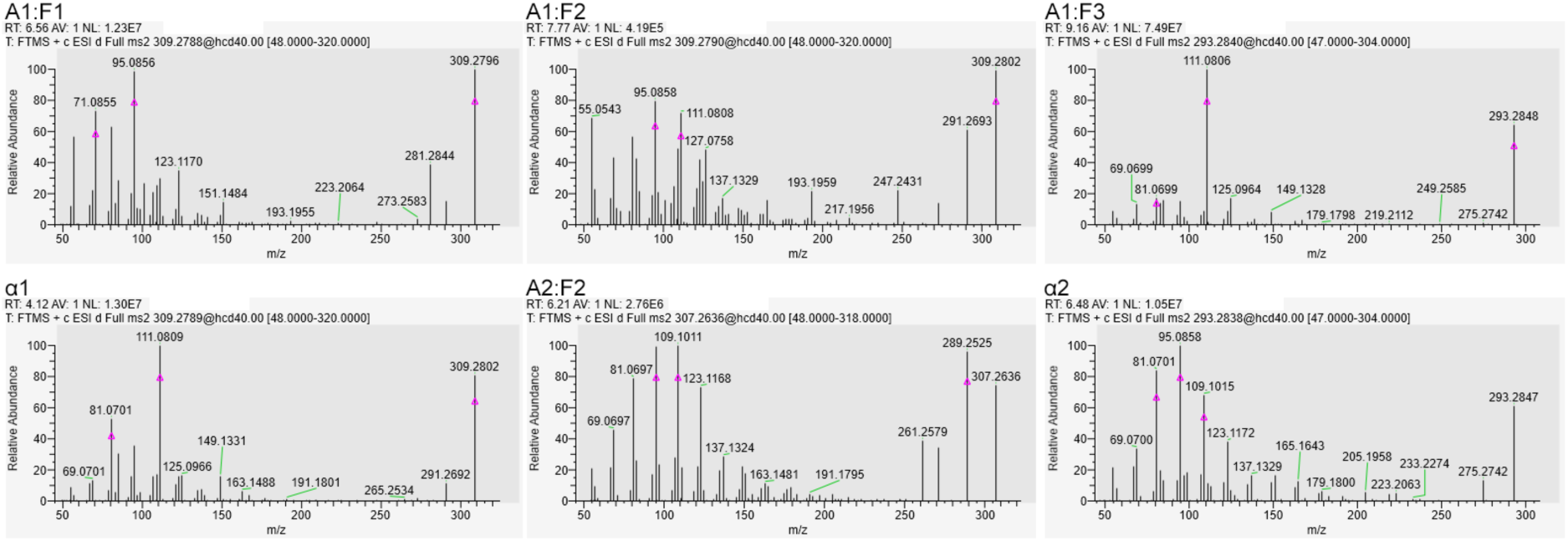
Fragmentation patterns (MS^2^) of three most abundant phytol metabolites in media extracts of A1 and A2 strains. Metabolite identity is shown above each plot on the first line. Second line shows retention time, (RT) and peak intensity (NL). Third line indicates acquisition parameters denoting FTMS detection in positive electrospray mode using data-dependent MS2 of the precursor ion at indicated m/z fragmented by HCD at 40% normalized collision energy, with a fragment scan range indicated in square brackets.

**Fig. S8.**
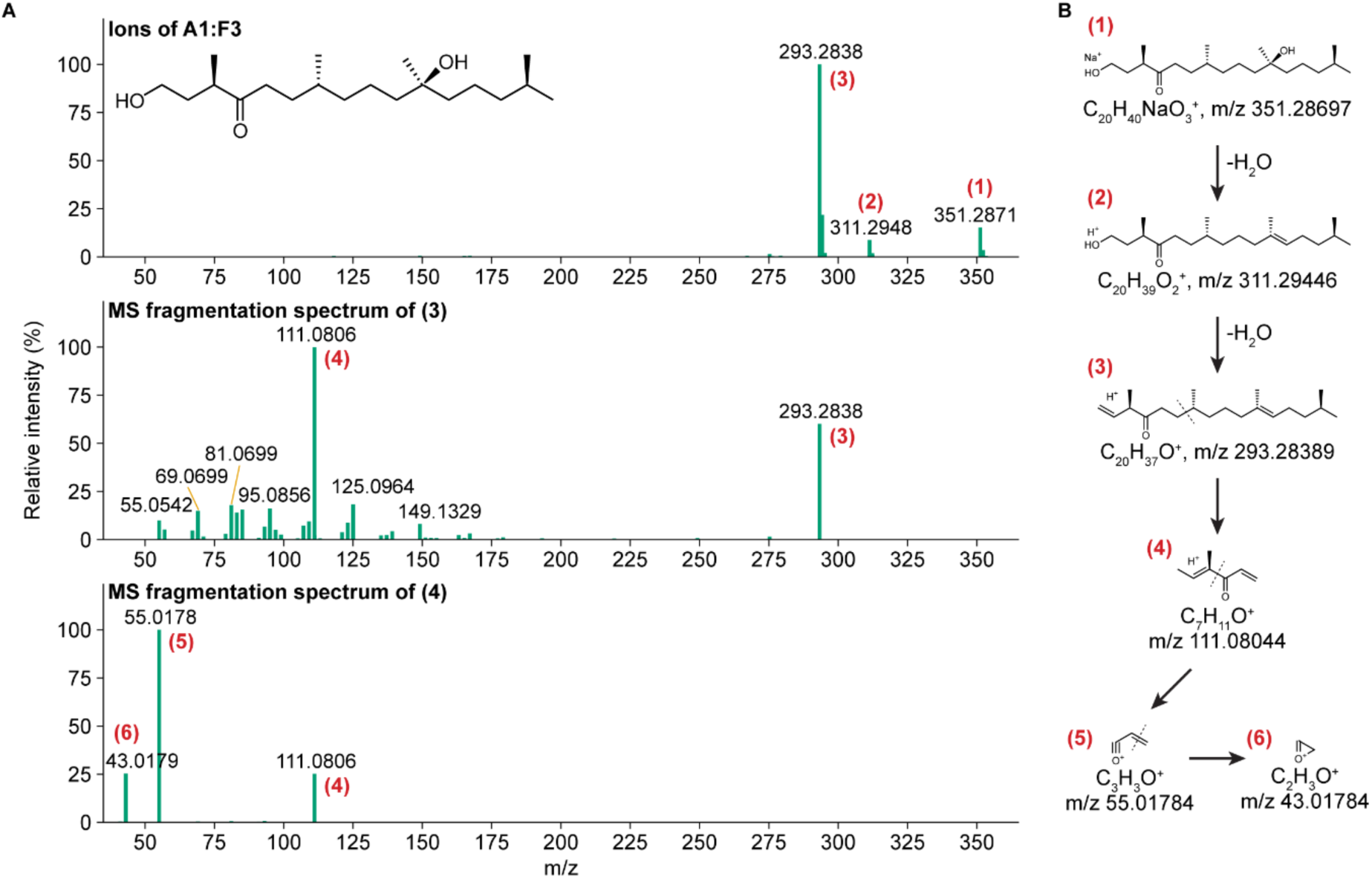
MS fragmentation analysis of the unlabeled A1:F3 metabolite. (A) Top: Detected ions of A1:F3 corresponding to (1) [M+Na]^+^, (2) [M-H2O+H]+ and (3) [M-2H2O+H]+. Middle: Fragmentation spectrum (MS^2^) of (3). Bottom: fragmentation spectrum (MS^3^) of (4). (B) Inferred structures of detected ions in the panel A, with broken C-C bonds points indicated by the dashed line.

**Fig. S9.**
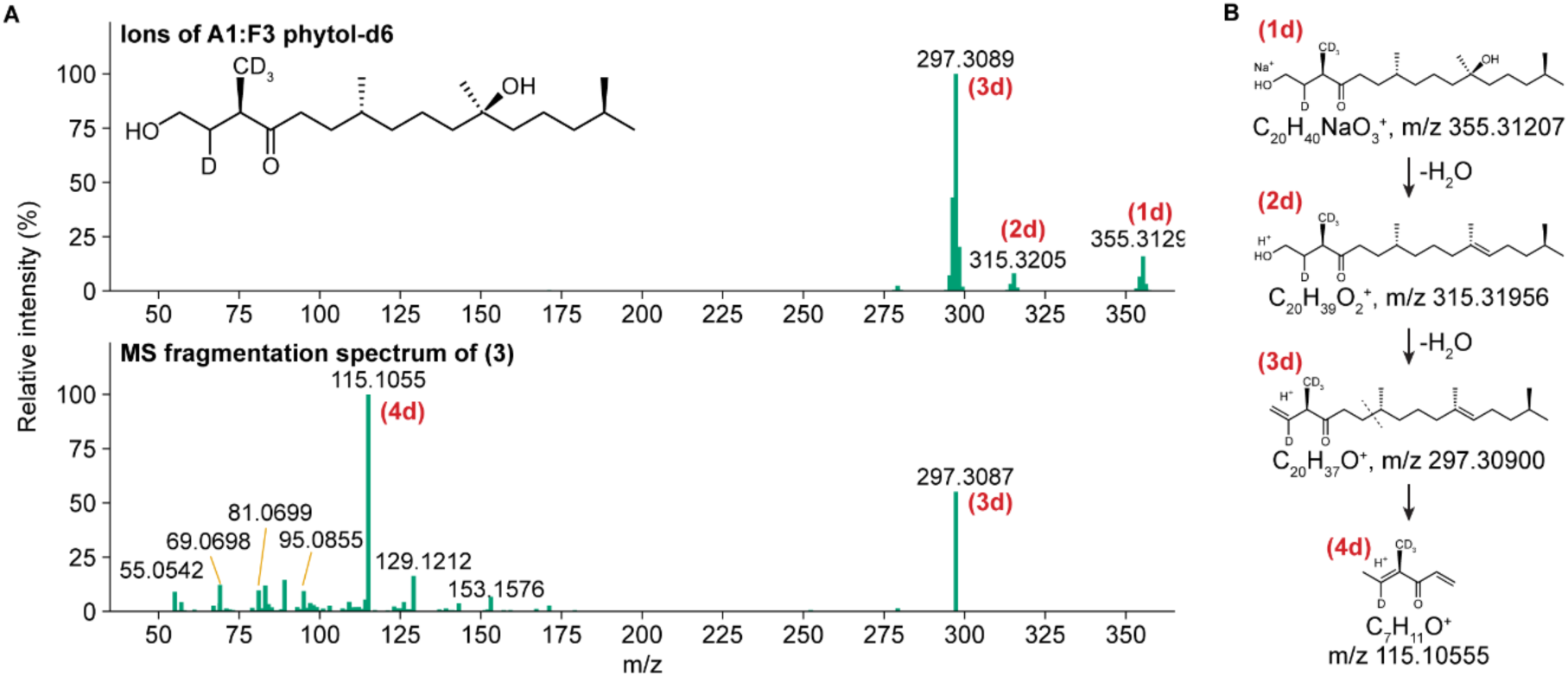
MS fragmentation analysis of the deuterium labeled A1:F3 metabolite, produced in phytol-d6 feeding experiment. (A) Top: Detected ions of A1:F3 corresponding to (1) [M+Na]^+^, (2) [M-H2O+H]+ and (3) [M-2H2O+H]+. Middle: Fragmentation spectrum (MS^2^) of (3). (B) Inferred structures of detected ions in the panel A, with broken C-C bonds indicated by the dashed line.

**Fig. S10.**
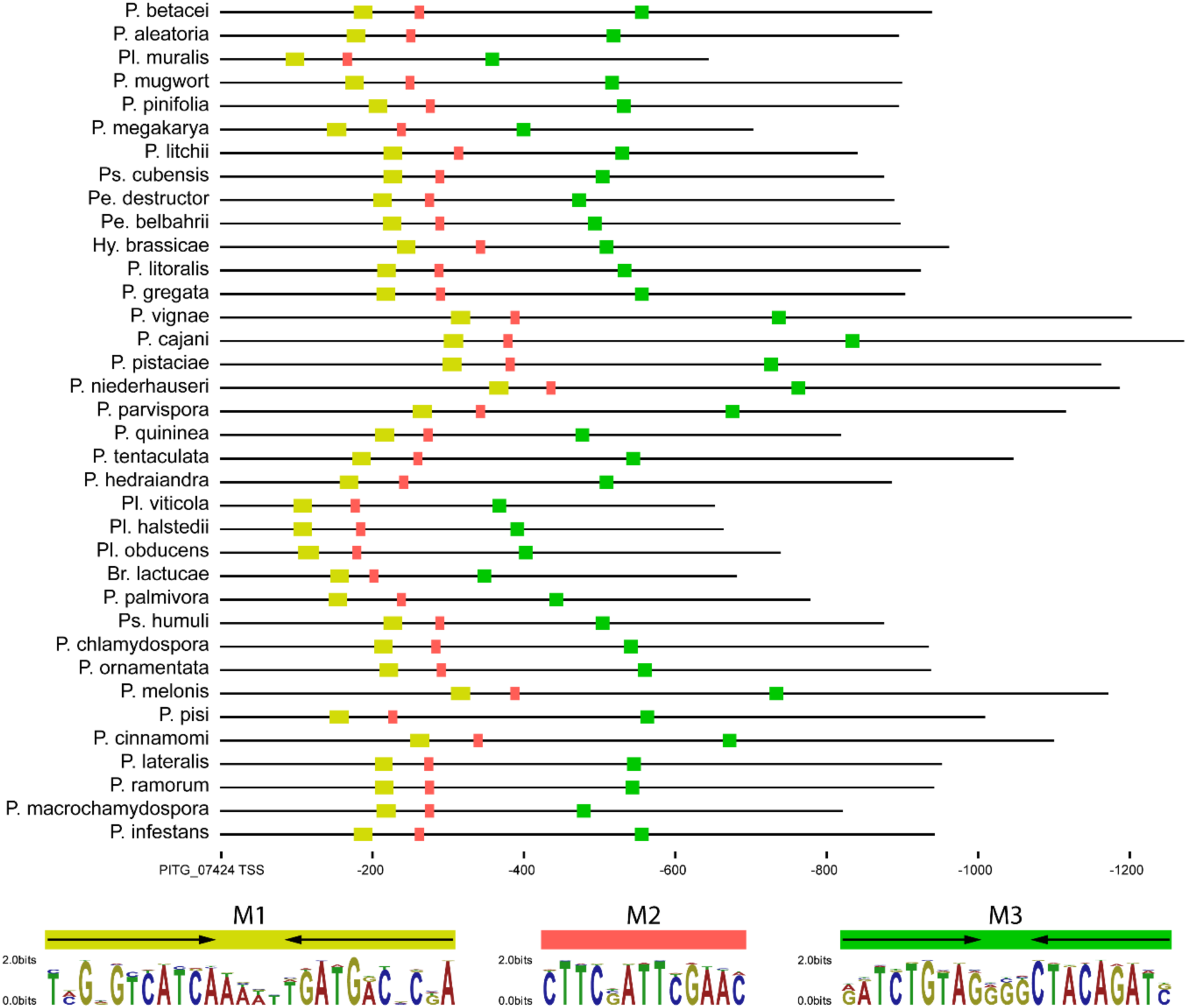
Positions of conserved motifs identified by phylogenetic footprinting across Peronosporales species in the bidirectional promoter region, between the transcription start sites of 07424 and 07423. Only species with conserved motifs are shown. Horizontal axis represents relative position from the PITG_07424 transcription start site motif. Colors represent different motifs. Corresponding motif sequence logos are shown at the bottom of the plot. Arrows indicate inverted within the motif.

**Fig. S11.**
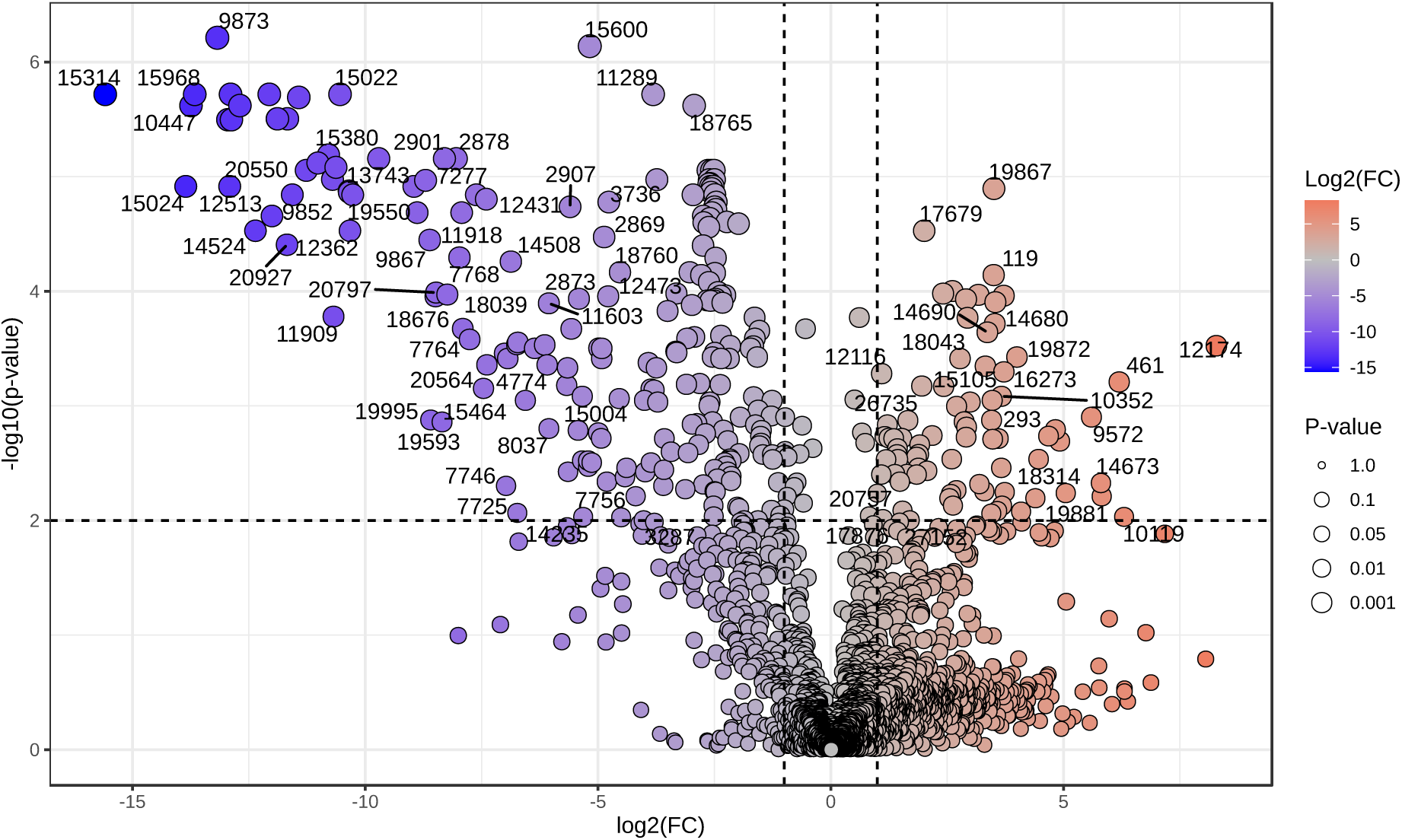
Volcano plot showing metabolite abundances in A2 strain following trichostatin A treatment.

**Fig. S12.**
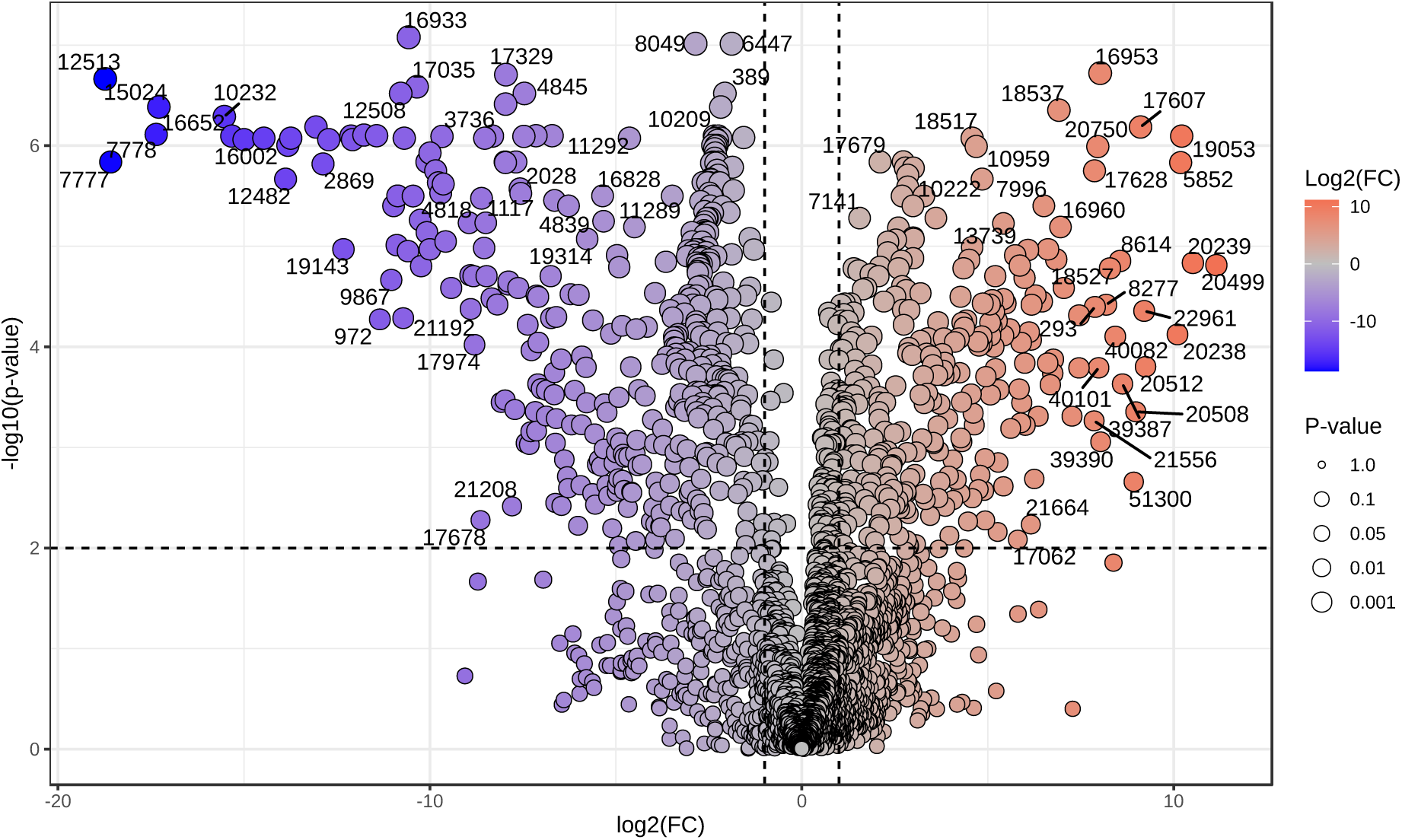
Volcano plot showing metabolite abundances in A1 strain following trichostatin A treatment.

**Fig. S13.**
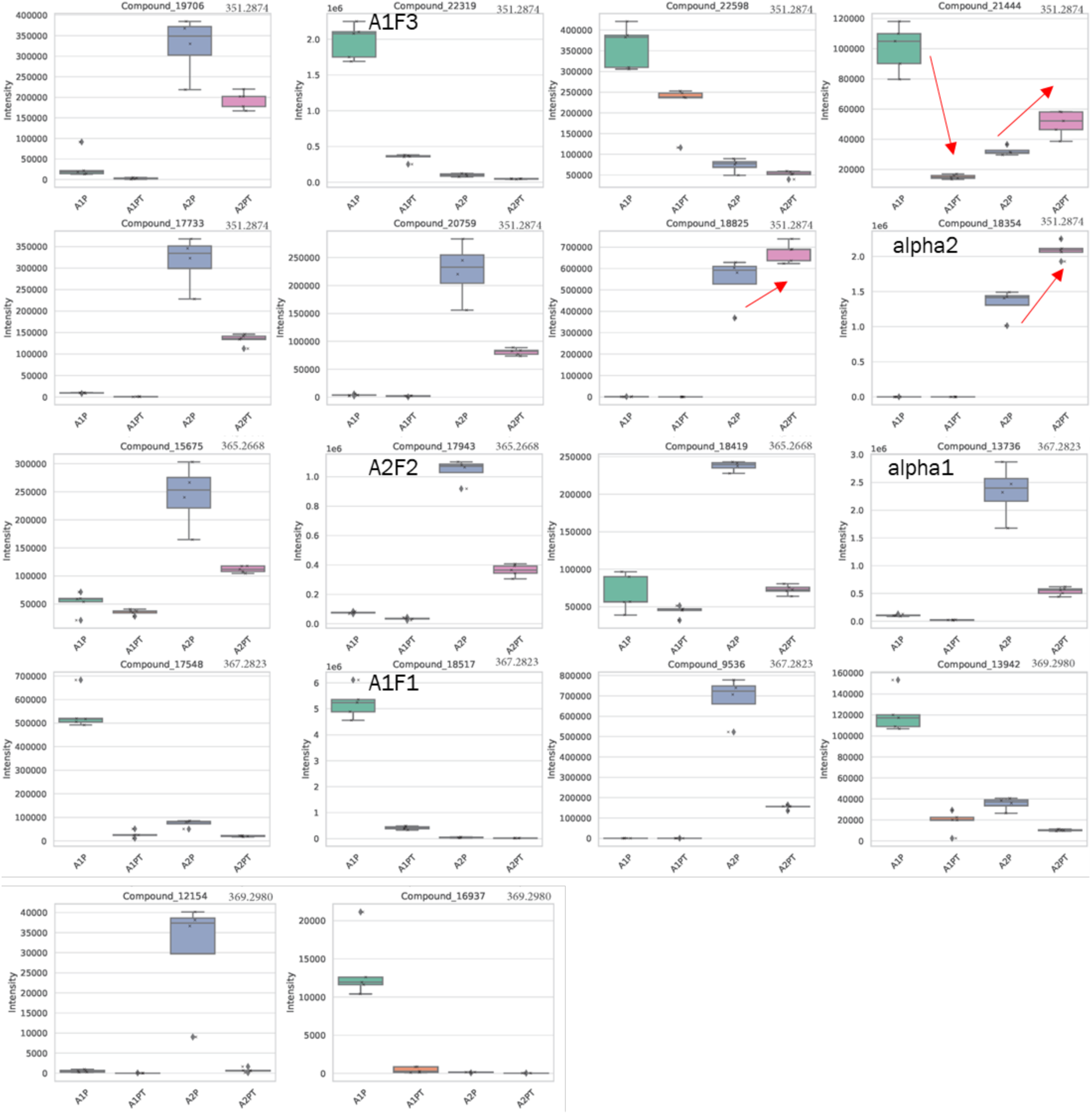
Change in phytol metabolites following trichostatin A (TSA) treatment in A1 and A2 strains. Major metabolites are annotated inside the plot area. Vertical axis represents intensity as the area under the peak. Where applicable, the axis is scaled by the value indicated in the upper left of the plot (1e6). Horizontal axis codes: A1P - A1 strain with phytol, A1PT - A1 strain with phytol and TSA, A2P - A2 strain with phytol, A2PT – A2 strain with phytol and TSA. Upper right of the plot is the m/z value. Red arrows indicate significant increases in intensity.

**Fig. S14.**
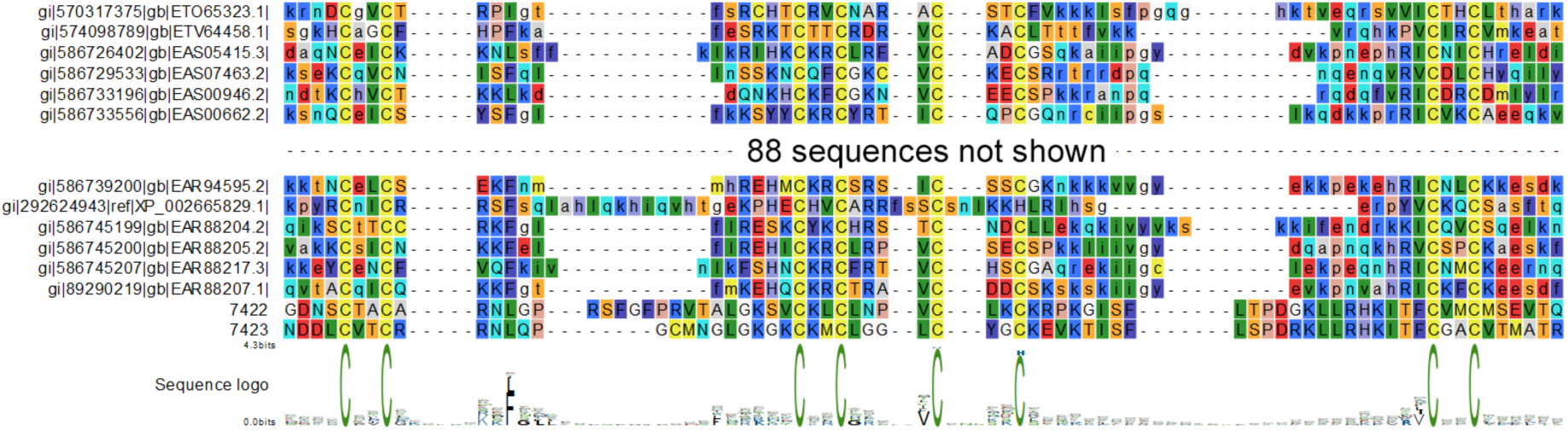
Protein sequence alignment showing FYVE domain of 0422, 07423 and 100 most diverse FYVE domain sequences from NCBI Conserved Domain database. Sequence logo at the bottom is derived from all sequences aligned, however 88 are omitted from the alignment display.

**Fig. S15.**
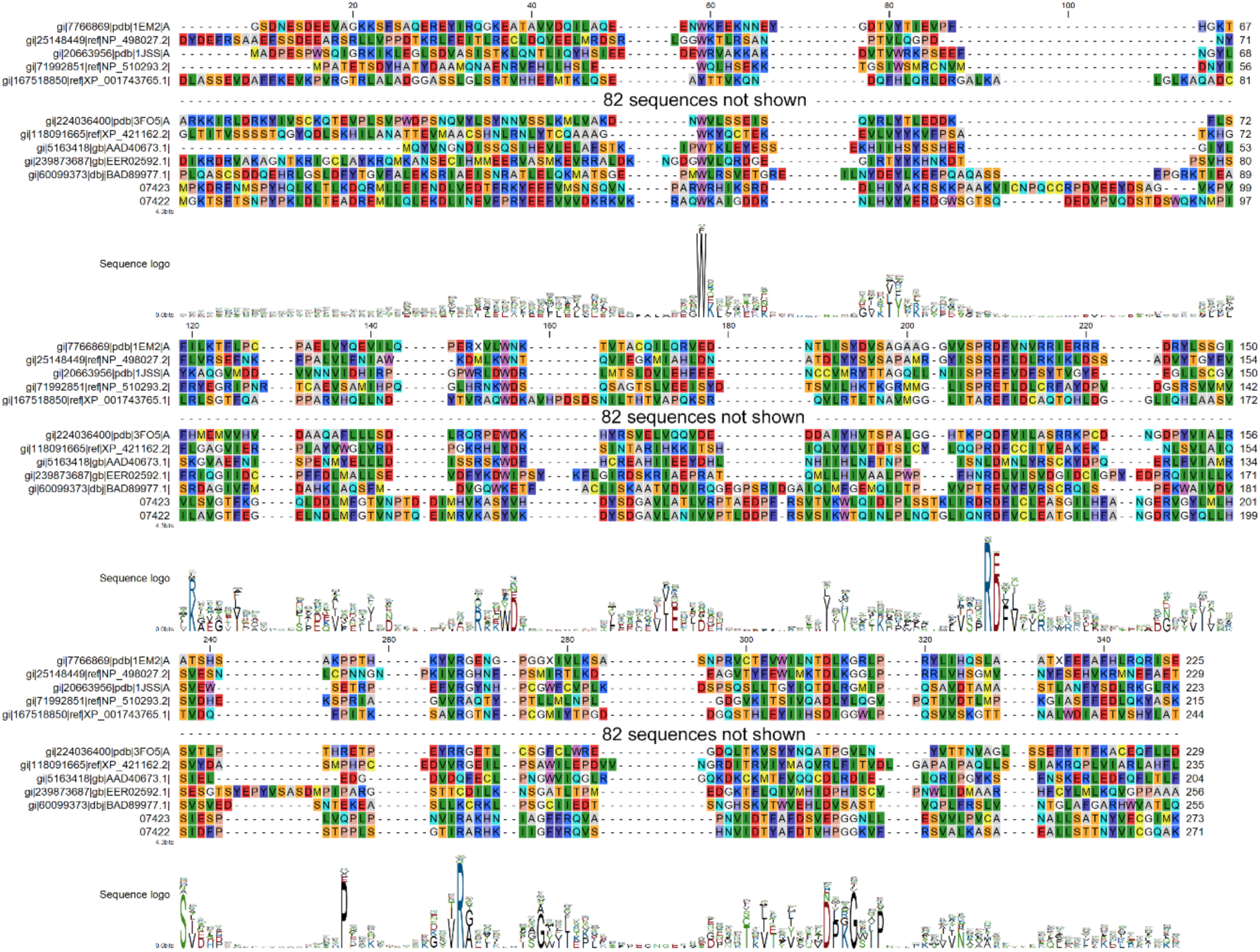
Protein sequence alignment showing START domain of 0422, 07423 and all (92) most diverse START domain sequences from NCBI Conserved Domain database. Sequence logo at the bottom is derived from all sequences aligned, however 82 are omitted from the alignment display.

**Fig. S16.**
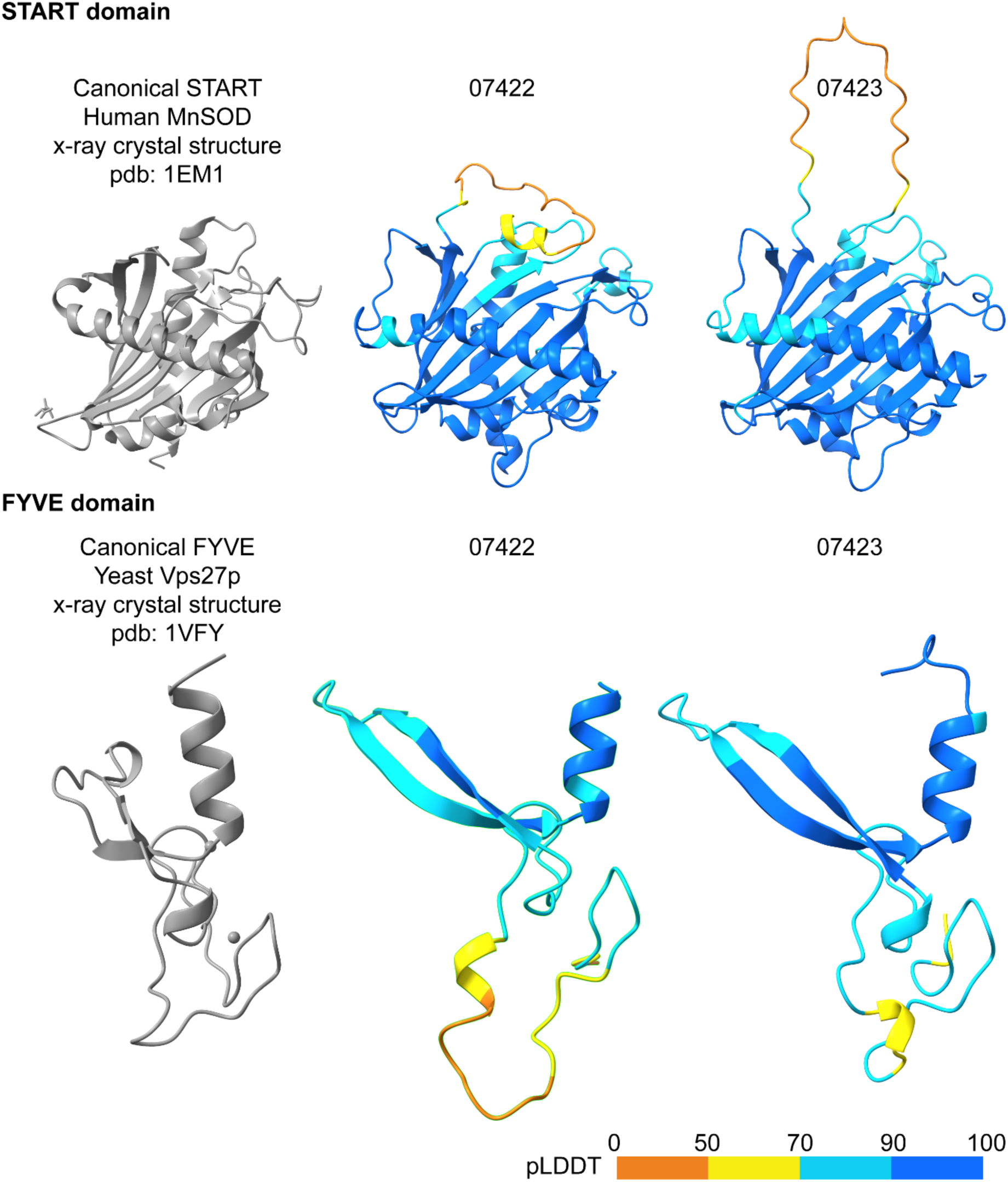
3D structure of canonical START (top) and FYVE (bottom) domains along with corresponding domains of 07422 and 07423. For both domains, structure of the canonical domains (shown in gray) are derived from x-ray crystallography and their identity is shown above the structure. Structures of 07422 and 07423 are predicted by AlphaFold3 and are colored by pLDDT (Predicted Local Distance Difference Test).

**Fig. S17.**
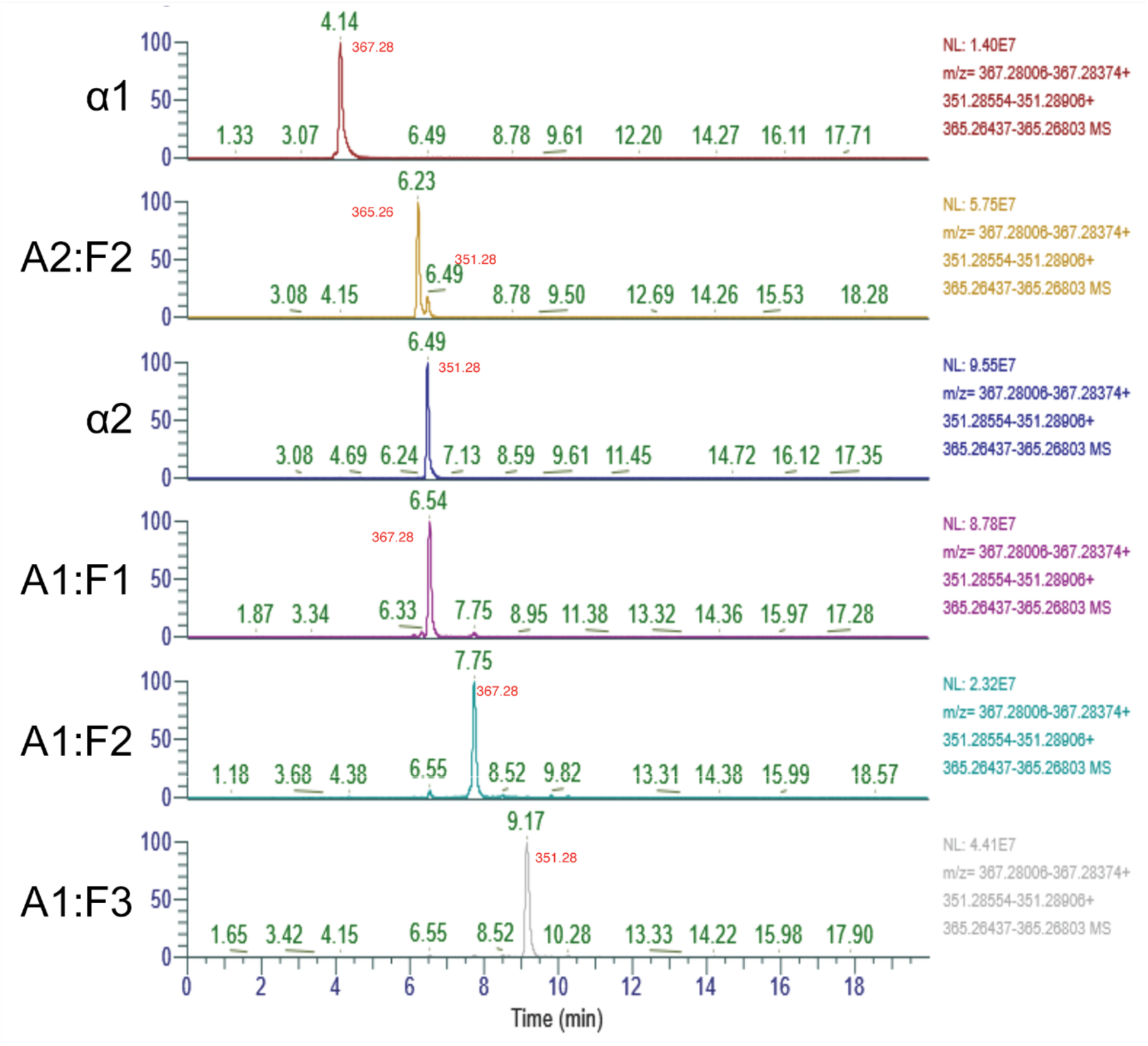
Extracted ion chromatograph (EIC) of *Phytophthora infestans* A1 and A2 media extracts after fractionation. Left of each chromatograph is the name of the tallest (target) peak. Green numbers above peaks show retention time. m/z value is shown in red for significant peaks. Absolute height of the tallest peak (NL) and extracted mass ranges (m/z) are shown to the right.

**Fig. S18.**
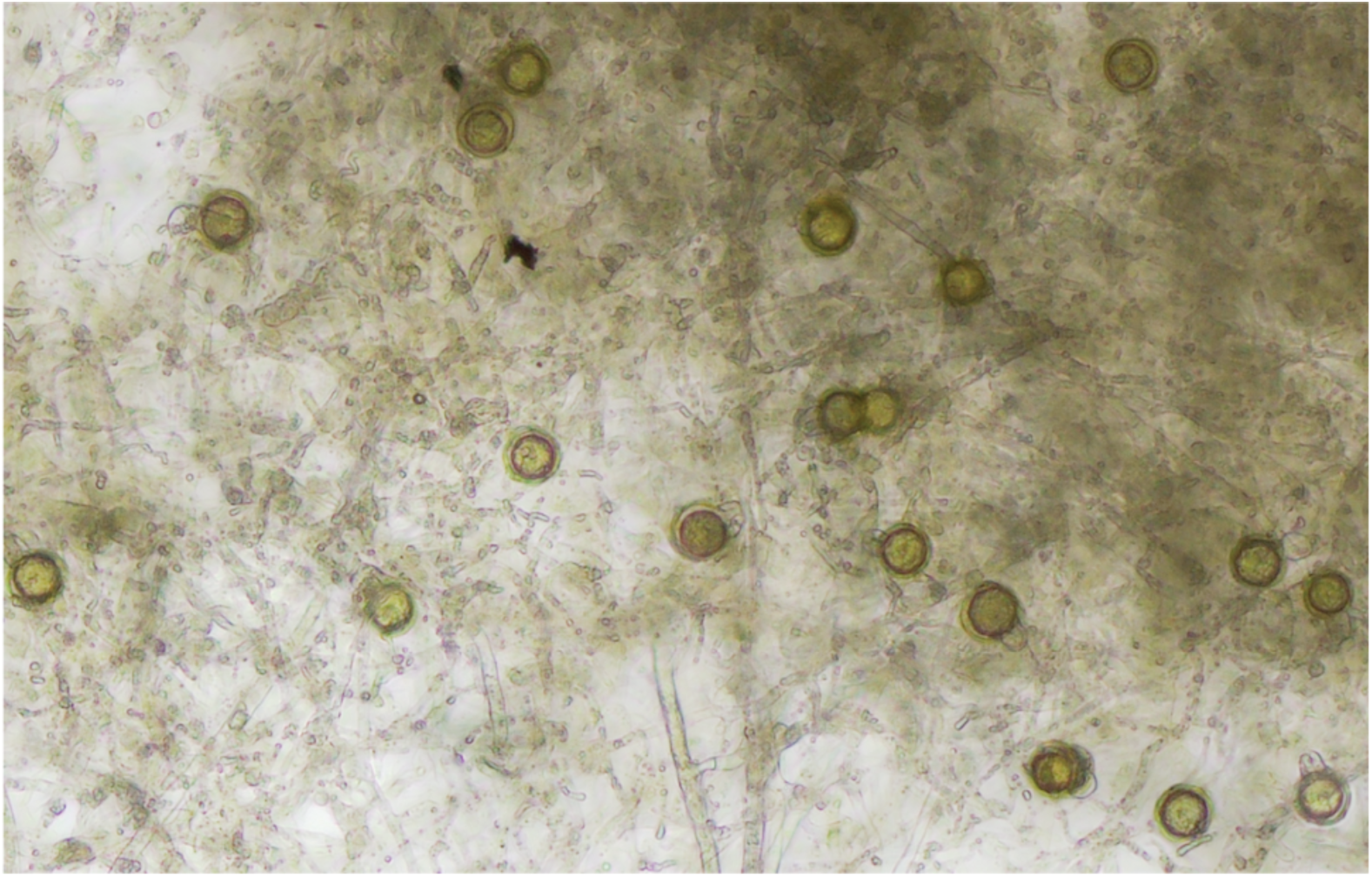
Bioassay demonstrating induction of oospore formation in an A1 strain of *Phytophthora nicotianae* following application of *Phytophthora infestans* α2 by paper-disc method.

**Table S1.** Tabular overview of cytochrome P5450 orthogroups across species, with copy number shown per species and per orthogroup.

**Table S2.** Metabolic conversion of phytol metabolite fractions by *Phytophthora infestans* A1 and A2 strains. Numbers represent area under the peak. Peak area of the original metabolite applied is bolded. Substantial conversion products are shown in red. Gray bar within each cell represents relative peak area within the treatment (column). D-atoms lost is shown for metabolites that were also detected in *P. infestans* cultures supplied with deuterium labeled phytol (phytol-d6, shown below the table). It represents number of deuterium atoms that were lost during phytol metabolism.

**Table S3.** Conservation of the mating cluster genes across Peronosporales species analyzed along with their mating system. For each species, HT signifies heterothallic, HO - homothallic, S - sterile, NA – unknown mating system. For orthologs of *P. infestans* PITG_07422, PITG_07423, and PITG_0724, the presence of an intact gene is indicated with the number 1, while damaged is indicated by 0.

**Table S4.** Oligonucleotide sequences used.

**Table S5.** IDs of BUSCO genes used in constructing accurate species-level phylogenetic tree.

**Table S6.** Accession numbers and metadata of genome assemblies used in the study.

**Table S7.** NCBI SRA run accession numbers of all raw read datasets used in the study along with corresponding metadata. Mating type assignments were based on the external literature.

